# The Interplay between Mutagenesis and Extrachromosomal DNA Shapes Urothelial Cancer Evolution

**DOI:** 10.1101/2023.05.07.538753

**Authors:** Duy D. Nguyen, William F. Hooper, Timothy R. Chu, Heather Geiger, Jennifer M. Shelton, Minita Shah, Zoe R. Goldstein, Lara Winterkorn, Michael Sigouros, Jyothi Manohar, Jenna Moyer, David Wilkes, Rahul R. Singh, Weisi Liu, Andrea Sboner, Scott T. Tagawa, David M. Nanus, Jones T. Nauseef, Cora N. Sternberg, Ana M. Molina, Douglas Scherr, Giorgio Inghirami, Juan Miguel Mosquera, Olivier Elemento, Nicolas Robine, Bishoy M. Faltas

**Affiliations:** Department of Medicine, Weill Cornell Medicine, New York, NY, USA; New York Genome Center, New York, NY, USA; Caryl and Israel Englander Institute for Precision Medicine, Weill Cornell Medicine, New York, NY, USA; Department of Pathology and Laboratory Medicine, Weill Cornell, New York, NY, USA; Sandra and Edward Meyer Cancer Center, Weill Cornell Medicine, New York, NY, USA; Department of Physiology and Biophysics, Weill Cornell Medicine, New York, NY, USA; Department of Urology, Weill Cornell Medicine, New York, NY, USA; Institute for Computational Biomedicine, Weill Cornell Medicine, New York, NY, USA; Department of Cell and Developmental Biology, Weill Cornell Medicine, New York, NY, USA

## Abstract

Advanced urothelial cancer is a frequently lethal disease characterized by marked genetic heterogeneity. In this study, we investigate the evolution of the genomic signatures caused by endogenous and external mutagenic stimuli and their interplay with complex structural variants. We superimposed mutational signatures and phylogenetic analyses of matched serial tumors from patients with urothelial cancer to define the evolutionary patterns of these processes. We show that APOBEC3-induced mutations are clonal and early, whereas mutational bursts comprising hundreds of late subclonal mutations are induced by chemotherapy. Using a novel genome graph computational paradigm, we observed frequent circular high copy-number amplicons characteristic of extrachromosomal DNA (ecDNA) involving double-minutes, breakage-fusion-bridge, and tyfonas events. We characterized the distinct temporal patterns of APOBEC3 mutations and chemotherapy-induced mutations within ecDNA, gaining new insights into the timing of these events relative to ecDNA biogenesis. Finally, we discovered that most *CCND1* amplifications in urothelial cancer arise within circular ecDNA amplicons. These *CCND1* ecDNA amplification events persisted and increased in complexity incorporating additional DNA segments potentially contributing selective fitness advantage to the evolution of treatment resistance. Our findings define fundamental mechanisms driving urothelial cancer evolution and have therapeutic implications for treating this disease.

## Introduction

During the lifetime of cancer, there is a constant interplay between mutagenesis and DNA repair that leaves behind detectable, distinct mutational signatures and extensive structural genomic changes^1–4^. Somatic mutational signatures characterized by cytosine to thymine or guanine substitutions that arise from expression of endogenous apolipoprotein B mRNA-editing enzyme catalytic subunit 3 (APOBEC3) of cytidine deaminases^5–7^ are commonly observed in the genomes of various human cancers, especially urothelial carcinoma (UC)^2, 5, 8^. In a previous study, we investigated the evolutionary dynamics of chemotherapy-resistant advanced UC^9^. However, several fundamental questions remain: (i) what are the relative timing, clonality, and velocity of endogenous mutagenic stresses such as APOBEC3 and extrinsic mutagenic processes such as chemotherapy in shaping urothelial cancer evolution? and (ii) what is the role of high-order structural variants, including extrachromosomal DNA (ecDNA), in urothelial cancer evolution and drug resistance? Without addressing these foundational questions, our ability to predict the clinical trajectory and intercept the development of drug resistance in patients with advanced cancer is limited.

To address these critical questions, we performed whole-genome sequencing of matched sets of primary, metastatic, and morphologically normal urothelial samples. We employed comprehensive analysis of single base substitution (SBS), doublet base substitution (DBS), and small insertions and deletions (ID) mutational signatures superimposed on phylogenetic analysis of primary and metastatic tumors collected at different time points. We compared the relative timing and clonality of mutagenic processes throughout the patients’ lifespans. We then applied a novel graph computational paradigm^10^ to serially collected tumor samples to analyze the topology of complex structural variants and investigate the evolution of ecDNA. We then integrated our in-depth mutational and structural variant analyses to characterize the patterns and timing of APOBEC3 and chemotherapy-induced mutations within ecDNA events. Finally, we mapped the changes in ecDNA events involving recurrent *CCND1* complex rearrangements following systemic therapy to gain insights into their role as putative drug-resistance mechanisms.

## Results

### Timing, clonality, and velocity of mutagenic processes shaping urothelial cancer evolution

We performed whole-genome sequencing of 77 urothelial tumors from 50 patients, including 27 pre-and 50 post-platinum chemotherapy samples, and characterized their mutational footprints. Since APOBEC3 cytidine deaminases are the predominant mutagenic enzymes in UC, we prioritized investigating SBS2^3, 11^, SBS13^3, 11^, and DBS11^12^ induced by APOBEC3. In addition, we also investigated SBS31^13^, SBS35^13^, and DBS5^2, 14^ associated with platinum-based chemotherapy, the backbone of the commonly used treatment regimen for advanced UC **(Supplementary Tables 1, 2 & Supplementary Figs. 1, 2)**. Across all samples, the mean contribution (percentage of all SBSs) of SBS2 was 17%, SBS13 was 20%, and the contribution (percentage of all DBSs) for DBS11 was 17% **(Supplementary Fig. 3a & Supplementary Table 2)**. The contribution of platinum-based chemotherapy-induced signatures SBS31 (mean 18%), SBS35 (mean 3%), and DBS5 (mean 23%) were dominant only in post-chemotherapy tumors **(Supplementary Fig. 3b & Supplementary Table 2)**.

To gain deeper insights into tumorigenesis and early tumor evolution, we established the timing and clonality of mutagenic processes across the history of tumor development by generating phylogenetic trees using variant cancer cell fraction (CCF) as input for 14 patients from whom we had at least two tumor samples. We then developed a method to superimpose the mutational signature composition onto the phylogenetic tree based on the variants of each branching point to illustrate dynamic changes in these signatures over time **(Methods, Fig. 1a, Supplementary Fig. 4a-m & Supplementary Table 3)**. In 11/14 reconstructed evolutionary trees, APOBEC3-induced signatures (SBS2/13) dominated the early truncal nodes comprising ≥ 25% of all mutations and persisted throughout the tumor’s lifetime **(Supplementary Fig. 4a-m & Supplementary Table 3)**, underscoring the role of APOBEC3 in initiating urothelial carcinogenesis and in its progression. In contrast, chemotherapy-induced mutations were late-appearing, mainly dominating the ‘branch’ nodes. This appearance is consistent with the relatively late timing of chemotherapy administration in the clinical course of the disease. We then used MutationTimeR^15^ to classify the clonality of somatic mutations in each of the 77 tumors **(Methods)**. We calculated for each tumor the enrichment of early variants (early-to-late clonal mutation fold change), and clonal variants (clonal-to-subclonal mutation fold change) contributed by mutational processes induced by APOBEC3 (SBS2/13) and chemotherapy (SBS31/35) **(Fig. 1b)**. Consistent with the pattern observed in the tumor phylogenies, APOBEC3-induced mutations were significantly earlier (1.85 versus 0.48-fold, *P* = 2.1×10^-7^) and more clonal (1.49 versus 0.8-fold, *P* = 4.3×10^-4^) compared to mutations attributed to platinum-based chemotherapy **(Fig. 1b)**. Next, we examined the mutational contribution over an estimated exposure time to estimate the mutagenic velocity of APOBEC3 and chemotherapy **(Methods, Fig. 1c & Supplementary Fig. 5)**. Platinum-based chemotherapy SBS31/35 and APOBEC3 SBS2/13 showed significantly higher mutagenic velocity compared to the natural aging signature SBS1 (*P* = 1.1×10^-19^ and *P* = 2.0×10^-19^ respectively) **(Fig. 1c)**. Importantly, chemotherapy exposure resulted in a higher mutational rate than did APOBEC3 for both SBS (SBS31/35 has 6.61-fold increase than SBS2/13, *P* = 9.4×10^-5^) and DBS-assigned mutations (DBS5 has 27.46-fold increase than DBS11, *P* = 6.0×10^-9^) even after accounting for different estimates for the onset of APOBEC3-induced mutagenesis **(Fig. 1c & Supplementary Fig. 5)**. Platinum-based chemotherapy resulted in a mean of 707.2 single-base substitutions and 3.9 double-base mutations per month of exposure **(Fig. 1c & Supplementary Fig. 5)**. These data show that chemotherapy induces extreme bursts of subclonal mutations even during relatively short time spans.

**Figure 1.**
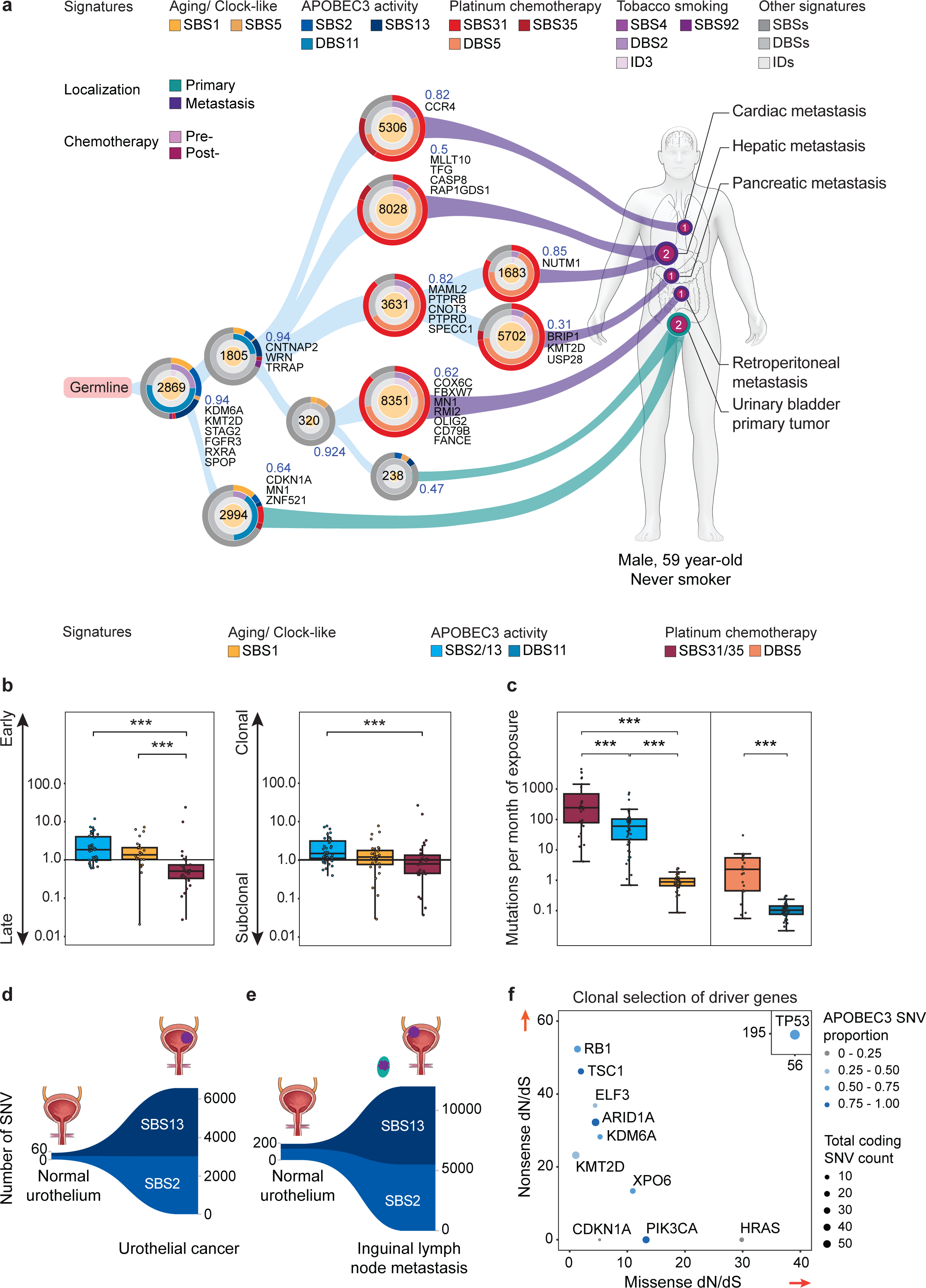
The relative timing of APOBEC3 and chemotherapy-induced mutagenesis in urothelial cancer evolution. **(a)** The superimposed mutational signatures and phylogenetic tree depict the evolutionary descent of tumor samples from a germline sample for patient WCMIV063. Each node represents the number of single nucleotide variants (SNVs) (centered number), the mean cancer cell fraction (CCF) across all SNVs in the node (blue number), and the high-impact SNVs in cancer census genes. In each node, SNVs were input into deConstructSigs to estimate the proportions of SBS, DBS, and small ID mutational signatures represented by three circle graphs from the periphery to the center, respectively. The tumor samples used for phylogenetic analysis are traced back to their corresponding anatomical sites on the schematic of the patient with annotation of their chemotherapy treatment status. The numbers on the human diagram represent the number of samples collected from the tumor sites. **(b)** The clonality fold-change of mutations contributed by combined SBS mutational signatures of APOBEC3 (SBS2/13), platinum chemotherapy (SBS31/35), and aging (SBS1) in the WCM-UC cohort. The early versus late clonality (left) and clonality versus subclonality (right) of the mutations were classified by MutationTimeR. Mutations contributed by platinum chemotherapy-associated signatures are enriched for late subclonal substitutions compared to signatures induced by APOBEC3, which are active earlier, or of aging, which is active throughout the lifetime of patients. Two-sided Wilcoxon rank sum test. ***: *P*<0.001. Boxplots show the median and IQR. The lower whisker indicates Q1-1.5*IQR. The upper whisker indicates Q3+1.5*IQR. Each dot represents one tumor. **(c)** Mutagenic velocity (average mutations per month of exposure) of platinum chemotherapy, APOBEC3 (10 years prior to the sample collection date), and aging-associated SBS and DBS signatures contributed to the mutational burden of tumors. Samples with no APOBEC3 signatures contribution and post-chemotherapy samples with no chemotherapy contribution were excluded. Two-sided t-test. ***: *P*<0.001. Boxplots show the median and IQR. The lower whisker indicates Q1-1.5*IQR. The upper whisker indicates Q3+1.5*IQR. Each dot represents one tumor. **(d, e)** Somatic mutations attributed to APOBEC3-induced signatures SBS2/13 were detected in morphologically normal urothelial samples of WCMIV001 and WCMIV013 and shared with their matched UC tumors (not indicated). Left and right y-axes represent the number of SNVs attributed to the signatures in a sample. **(f)** Scatter plot showing the maximum likelihood estimates of dN/dS ratios for all significant genes (q-value < 0.1). The size of each point reflects the number of coding mutations detected across the WCM-UC cohort for a given gene, and the color reflects the proportion of coding mutations attributable to APOBEC3 mutagenesis. *TP53* had estimated dN/dS values much higher than the other significant genes and is depicted in an inset plot in the top-right corner for visual clarity. dN/dS = 1: neutral selection, dN/dS > 1: positive selection, dN/dS < 1: negative selection. Higher missense dN/dS indicates clonal selection of oncogenes. Higher nonsense dN/dS indicates clonal selection of tumor suppressor genes. Arrows indicate the direction of increased clonal selection.

Based on the early and clonal distribution of APOBEC3-induced single base and doublet base substitutions **(Fig. 1b)**, we hypothesized that this process contributes to driving urothelial carcinogenesis. We identified hundreds of APOBEC3-induced mutations in morphologically normal urothelial tissue samples shared with urothelial cancers from the same patients **(Fig. 1d, e & Supplementary Table 4)**. Next, we analyzed the ratio of nonsynonymous mutations to synonymous mutations (dN/dS) in the UC tumors in WCM-UC cohort^16–18^ to identify APOBEC3-induced mutated genes that were clonally selected, indicating a potential fitness advantage probably started as early as the normal or pre-malignant stages of tumor development. This method of calculating the dN/dS ratio adjusts for other confounders of clonal selection, such as large gene sizes and a high regional mutation rate^19^. We identified an enrichment of APOBEC3-induced mutations in critical oncogenes (*PIK3CA*) and tumor suppressor genes (*TP53*, *RB1, KDM6A, ARID1A*) with dN/dS > 10, suggesting that APOBEC3-induced mutations in these genes have undergone intense and persistent selection, potentially providing a fitness advantage that extends back to before tumorigenesis **(Fig. 1f & Supplementary Table 5)**. Furthermore, consistent with the processive nature of APOBEC3 enzymes which can mutate successive cytidines on a single strand of DNA, we identified an enrichment of SBS2/13-induced composite mutations^20^ in multiple cancer-associated genes (*TP53, KMT2D, ELF3, CSMD3, KDM6A, KMT2A, MUC16, ARID1A, ARID2, NIN, PIK3CA, RB1, TPR)* **(Supplementary Table 6)**. These additional mutations in the same gene can potentially augment the selective fitness of the initial mutations^20^, prompting urothelial carcinogenesis. Our data are consistent with the emerging evidence of APOBEC3-induced mutations in histologically normal urothelium from organ transplant donors^21^.

Our findings support a paradigm in which APOBEC3-induced mutagenesis acts as an early driver of urothelial carcinogenesis. This is particularly significant given that APOBEC3-induced mutagenesis accounts for 67% of all SNVs in bladder cancer^22^. Furthermore, our data show the interplay between the early clonal APOBEC3-induced mutations and the late subclonal bursts of chemotherapy-induced mutations. These two forces cooperate to ultimately shape the evolution of the metastatic and treatment-resistant phenotypes.

### Complex structural variants, including extrachromosomal DNA in urothelial cancer

Genomic instability involves the interplay between mutations and complex, large-scale chromosomal variants. To investigate these processes, we utilized the Junction Balance Analysis (JaBbA)^10^ framework, which analyzes junctions between genomic intervals that are not contiguous in the reference genome^23^ to provide a comprehensive view of higher-order structural variants (SVs) consisting of multiple breakpoints with high copy number (CN) states in our WCM-UC cohort **(Methods, Fig. 2a, Supplementary Table 7 & SV Glossary)**. We identified a median of 19 SVs per sample (IQR: 9 – 36); 57.8% of the samples had two or more complex SVs, 84.5% had one or more complex SVs (median: 2, IQR: 1 – 5), and 15.5% had none **(Fig. 2a & Supplementary Table 7)**. Templated insertion chains (TIC) (49.3%), chromoplexy (33.8%), and breakage-fusion-bridges (BFBs) (31%) were the most prevalent complex SV events in our WCM-UC cohort **(Supplementary Fig. 6a & Supplementary Table 7)**. Interestingly, 11.3% of UC tumors harbored nested accumulation of junctions with high-JCN (junction copy number) and fold-back inversion events designated as “typhonas” by JaBbA^10^. These events carried a significantly higher SV junction burden compared to other complex SVs (*P* < 7.8×10^-5^, Wilcoxon rank-sum test) **(Supplementary Fig. 6b & Supplementary Table 7)**.

**Figure 2.**
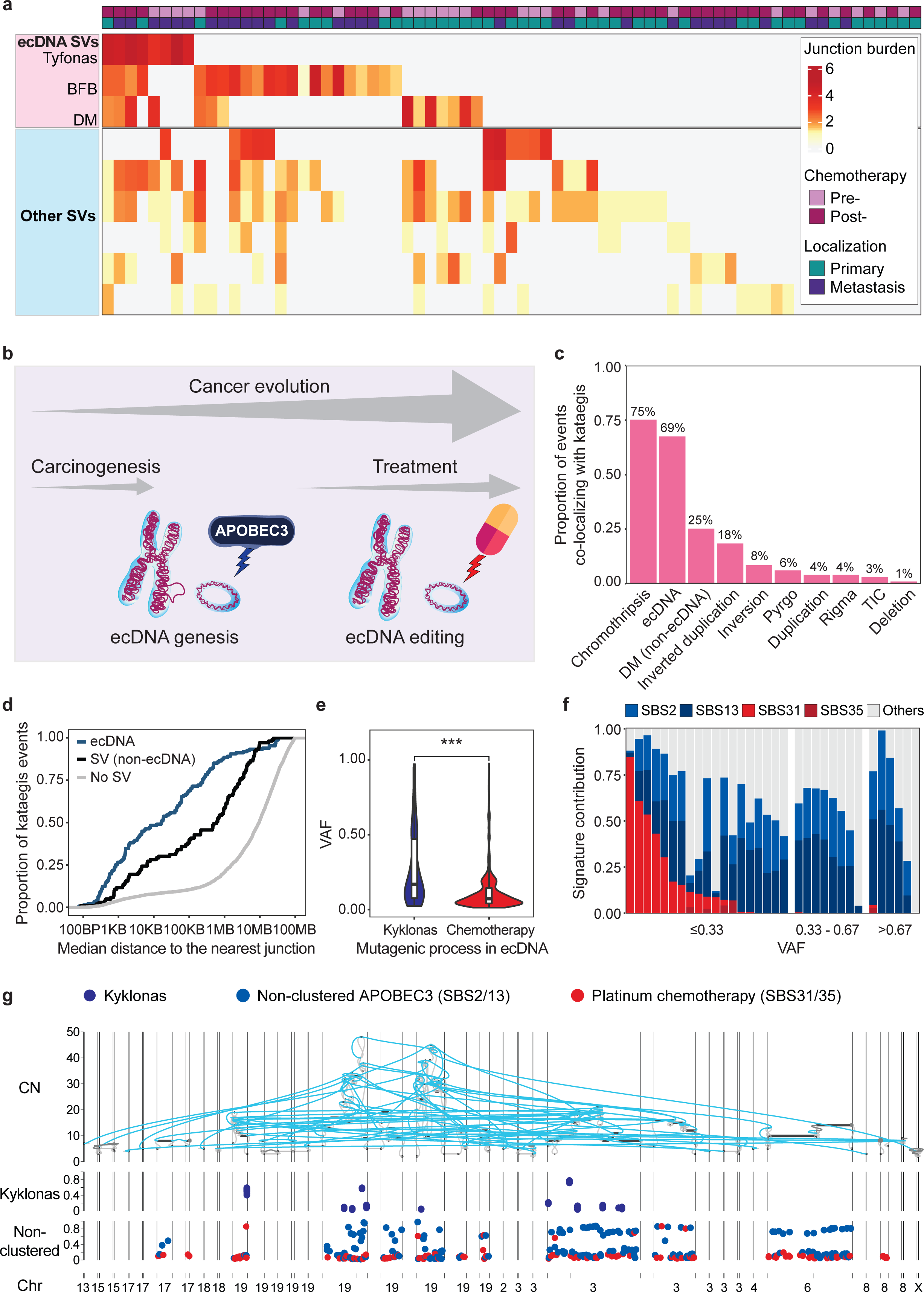
The interplay between APOBEC3 and platinum-chemotherapy induced mutagenesis and extrachromosomal DNA during urothelial cancer evolution. **(a)** A heatmap of the junction burdens of complex structural variants called by JaBbA (y-axis) in 71 UC tumors (x-axis). Tyfonas, breakage-fusion-bridges (BFBs), and double minutes (DMs) are grouped by their potential mechanisms to produce cyclic extrachromosomal DNA (ecDNA). The junction burden heatmap is normalized to the average junction burden of the WCM-UC cohort and scaled in the natural log. Other SVs: other complex SVs, from top-bottom: chromothripsis, chromoplexy, templated-insertion chains, quasi-reciprocal pairs, rigma, and pyrgo. **(b)** A schematic of the interplay between APOBEC3 and chemotherapy-induced mutageneses on ecDNA from its biogenesis and throughout cancer’s lifetime. **(c)** The proportion of all detected kataegic events co-localizing with structural variants footprints. The DM (non-ecDNA) class indicates double minutes identified by JaBbA but not classified as ‘cyclic’ by AmpliconArchitect. TIC: templated-insertion chains. **(d)** Kataegic events on ecDNA (teal) harbored a significantly shorter median distance to the nearest breakpoint than kataegis on other non-ecDNA structural variants (black) and kataegis with no SV association (gray). The median size of ecDNA in our WCM-UC cohort was approximately 5.56 Mb, so there was an upper limit of the distance to the closest breakpoints for kyklonas. The median distance of all mutations within a kataegic event collapsed to one measurement per event. *P* < 0.001, two-sided Wilcoxon rank sum test. **(e)** Variant allele frequency (VAF) distributions for APOBEC3-induced kyklonic mutations and chemotherapy-induced somatic mutations on ecDNA. ***: *P* < 0.001, two-sided Wilcoxon rank sum test. Violin plots extend between the maximum and minimum of the distribution. The middle boxplots show the median with IQR. The lower whisker indicates Q1-1.5*IQR. The upper whisker indicates Q3+1.5*IQR. **(f)** Mutational signature contributions to each sample with ecDNA/VAF tranche combination indicate the early APOBEC3 mutagenesis on ecDNA. Only sample/VAF tranche combinations with greater than 100 mutations are displayed. **(g)** The graph shows an ecDNA event harboring at least one kyklonic event in WCMIVG035S01. The top “CN” track represents the JaBbA genome graph showing copy-number alterations for rearranged DNA segments (gray vertices) with SV junctions (aqua blue edges) that form the circular ecDNA events in the urothelial tumor sample. The middle “Kyklonas” track shows the normalized VAF of identified APOBEC3-induced kyklonas. The bottom “Non-clustered” track shows normalized VAF of non-clustered mutations assigned to both APOBEC3-associated (SBS2/13) and platinum chemotherapy-induced (SBS31/35) mutational signatures on ecDNA. Chr: chromosome.

Tumors harboring *TP53* high and moderate impact mutations exhibited a significantly higher fraction of the genome altered (*P* < 0.001, Wilcoxon rank-sum test) and a significant increase in the number of total junctions (*P* = 4.9×10^-7^), deletions (*P* = 2.8×10^-3^), duplications (*P* = 2.0×10^-7^), chromoplexy (*P* = 3.6×10^-3^), TIC (*P* = 8.6×10^-3^), and BFB (*P* = 4.6×10^-5^) **(Supplementary Fig. 7)**. These findings suggest that background genomic instability due to mutations in guardians of genomic integrity, such as *TP53,* plays a critical role in the biogenesis of complex SVs.

Cancer cell ecDNA is a subclass of circular DNA in the nucleus characterized by large size (>1 Mb) and high-copy number amplification of oncogenes^24^. The biogenesis of ecDNA involves several potential mechanisms, including double minutes (DMs) and BFBs^10, 25^ complex SVs. Additionally, ecDNA results from catastrophic DNA breakage and ligation events^26^, such as chromothripsis^27^ and tyfonas^10^. As tyfonas, BFBs, and DMs are prevalent in our WCM-UC cohort and have been implicated in the biogenesis of ecDNA^10, 28–31^, we directly investigated the contribution of ecDNA to UC cancer evolution **(Fig. 2a)**. To examine the hypothesis of JaBbA’s tyfonas/BFBs/DMs to produce ecDNA events, we used a method specifically developed to reconstruct the circular configuration of focal amplification called AmpliconArchitect^32^ **(Methods)**. We found that tyfonas (100%), BFBs (81.5%), and DMs (63.2%) were the three most common JaBbA events to overlap with AmpliconArchitect’s cyclic event calls, supporting their role in ecDNA formation^10, 25, 33^ **(Supplementary Fig. 8)**. We discovered that 35% of samples in our WCM-UC cohort harbored one or more ecDNA event, demonstrating the high prevalence of ecDNA in UC.

### The interplay between APOBEC3 and chemotherapy-induced mutagenesis and extrachromosomal DNA during urothelial cancer evolution

We reasoned that the juxtaposition of APOBEC3 and chemotherapy-induced mutations on ecDNA could provide insights into the timing and mechanisms of ecDNA biogenesis **(Fig. 2b)**. Among 42 ecDNA events detected in our WCM-UC cohort, 69% showed evidence of kyklonas, defined as APOBEC3-induced clustered mutations (kataegis) occurring in ecDNA^34^ **(Fig. 2c, Supplementary Fig. 9a-i & Supplementary Table 8).** Kyklonic events occurred near SV breakpoints with a median distance of 16.2 KB between kyklonic events and the nearest breakpoint. 48.6% of kyklonic events occurred within 10 KB of an SV breakpoint **(Fig. 2d & Supplementary Table 8)**. Kyklonas were significantly closer to SV junctions compared to chromosomal kataegic events (*P* < 0.001, two-sided Wilcoxon rank sum test) **(Fig. 2d & Supplementary Table 8)**. The topographic proximity between APOBEC3-induced kyklonic events and SV breakpoints suggests a potential role for APOBEC3-induced DNA breaks as an intermediary step in the process of ecDNA biogenesis.

To investigate the relative timing of APOBEC3 and chemotherapy-induced mutagenesis in relation to ecDNA biogenesis, we examined the variant allele frequency (VAF) of APOBEC3 kyklonic clusters and non-clustered chemotherapy mutations on ecDNA. The average VAF of kyklonas was significantly higher than ecDNA-associated chemotherapy mutations (*P* < 2.22 × 10^-16^, two-sided Wilcoxon rank sum test) **(Fig. 2e & Supplementary Table 8)**. 10.2% of kyklonic mutations occurred early in the evolution of the ecDNA population among urothelial tumors (VAF > 0.67) **(Fig. 2f, g, Supplementary Fig. 9a-i & Supplementary Table 8)**. Notably, 93.4% (669/716) of somatic mutations induced by chemotherapy on ecDNA had low VAF < 0.33 **(Fig. 2f, g, Supplementary Fig. 9a-i & Supplementary Table 8)**. Together, these findings point to topographical and chronological overlap between the timing of APOBEC3-induced mutagenesis and ecDNA biogenesis.

### Recurrent extrachromosomal DNA-driven oncogene amplification in urothelial cancer

To identify recurrent extrachromosomal DNA-driven oncogene amplification, we searched for cancer-related genes near significantly recurrent breakpoints (SRB) –regions of recurrent structural variation where DNA can be inserted, deleted, or rearranged to form complex SVs. We applied the FishHook^35^ tool to search for genomic regions harboring these significantly recurrent breakpoints **(Methods)**. FishHook identified 31 loci, 17 (55%) of which were within 500 kb of a cancer-related gene (*CCND1, AXIN2, CHEK2, ERBB2, FHIT, KRAS, MDM2, NUMA1*), and 8/17 of these were less than 100 kb from a cancer-related gene (*CCND1, CHEK2, ERBB2, FHIT, NUMA1*) **(Fig. 3a & Supplementary Table 9)**. We then overlapped JaBbA SV classes with FishHook hits to further characterize the nature of the SVs involving these significantly recurrent breakpoints. Interestingly, the 11q13.3 locus harboring *CCND1* was involved in the highest number of SVs (13 samples), 92% of which were ecDNA-associated DM, BFB, and tyfonas events **(Fig. 3b & Supplementary Table 9)**. These events had a mean CN of 47 and a 3.6-fold higher mean CN than other non-cyclic amplifications of *CCND1* **(Fig. 3b & Supplementary Table 10)**.

**Figure 3.**
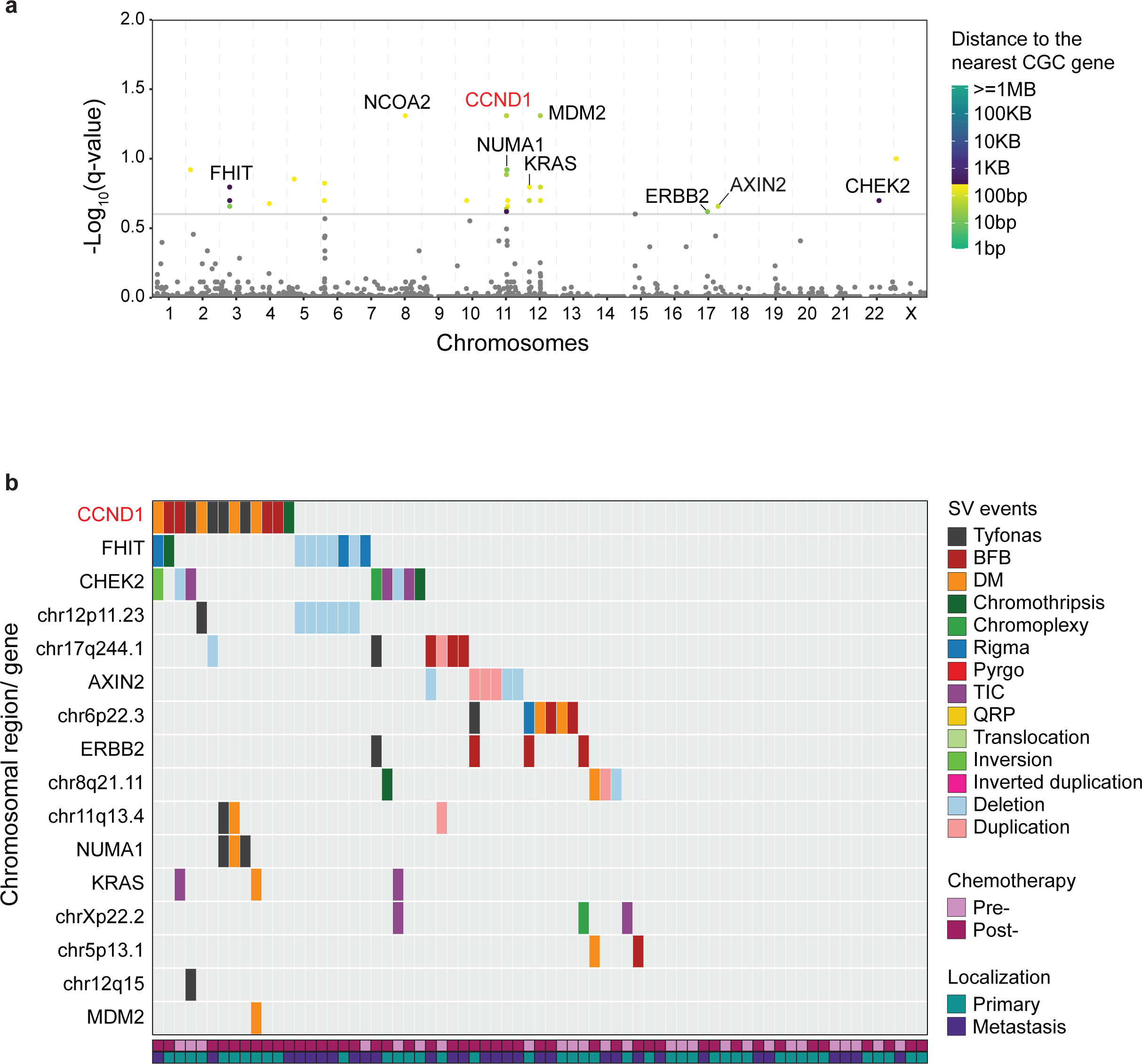
*CCND1* is commonly involved in recurrent putative ecDNA complex SVs in UC. **(a)** Manhattan plot shows the significantly recurrent breakpoints (SRBs) identified by FishHook and their distance to the nearest Cancer Gene Consensus (CGC) genes in UC whole-genome sequences. Each dot represents an FDR-adjusted (Benjamini-Hochberg) p-value of the distance, and a cutoff of 0.25 (horizontal solid line) was used to nominate significant hits. **(b)** JaBbA SV events were overlapped with FishHook SRB hits to identify the frequency and class of SV events occurring in significantly recurrent breakpoint regions with the nearest CGC genes (left y-axis) in each tumor (x-axis). The panel was arranged by decreasing the total number of SV events in a particular chromosomal region.

### Extrachromosomal DNA is the critical mechanism of *CCND1* amplification in urothelial cancer and a putative driver of resistance

Using AmpliconArchitect and JaBbA, we identified ecDNA events in 67% (10/15) of tumors harboring *CCND1* amplification in our WCM-UC cohort **(Supplementary Table 10)**. To further validate these findings, we examined *CCND1* ecDNA amplification in The Cancer Genome Atlas (n=3,731) and Pan-Cancer Analysis of Whole Genomes (n=1,291) (TCGA/PCAWG) pan-cancer cohorts by analyzing the AmpliconArchitect dataset^4, 36^. The TCGA/PCAWG Bladder Cancer cohort (BLCA-TCGA) harbored the highest proportion of circular ecDNA-based *CCND1* amplification (80%), followed by pancreatic adenocarcinoma (75%), ovarian carcinoma (67%), and esophageal carcinoma (59%) **(Fig. 4a & Supplementary Table 11)**. These data confirm that ecDNA formation is the dominant mechanism underlying *CCDN1* amplification in UC.

**Figure 4.**
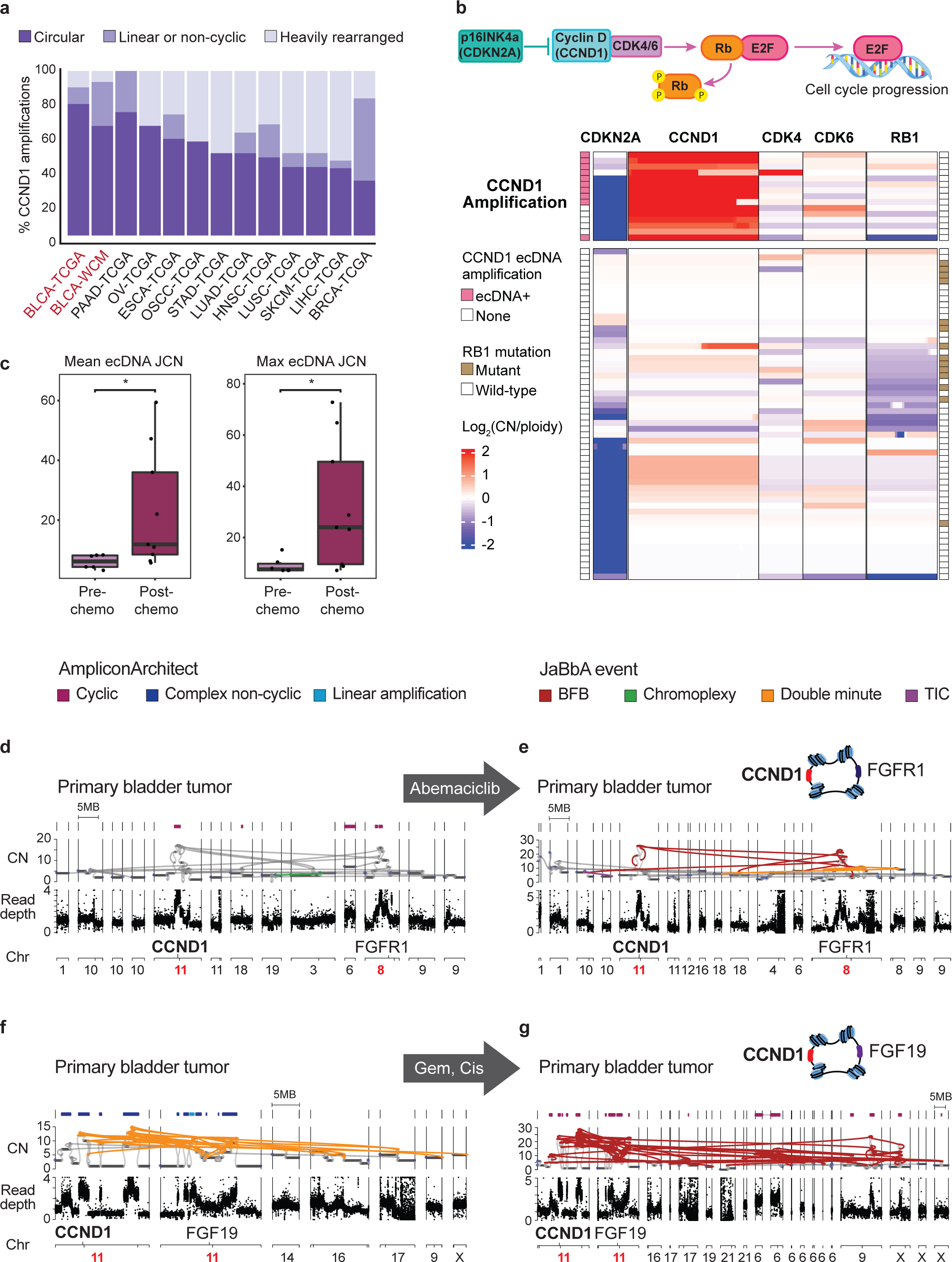
The increase in complexity and copy-number of ecDNA *CCND1* amplicons following systemic therapy of urothelial cancers. **(a)** Distribution of circular, heavily rearranged, linear and non-cyclic amplification mechanisms of focal somatic copy number amplification of *CCND1* across 71 samples from our Weill Cornell Medicine bladder cancer (BLCA-WCM) cohort and whole genomes from the TCGA/PCAWG pan-cancer cohorts as analyzed in *Kim et al. 2020*^36^. The study abbreviations for different cancer types are listed at https://gdc.cancer.gov/resources-tcga-users/tcga-code-tables/tcga-study-abbreviations. **(b)** (Top) A schematic of the p16-cyclinD1-CDK4/6-Rb pathway, a master regulatory mechanism of cell cycle progression. (Bottom) The normalized copy-number alteration heatmap of gene-containing chromosomal regions 9p21.3 (*CDKN2A*), 11q13.3 (*CCND1*), and genes *CDK4/6* and *RB1* in 71 UC tumor samples. The size of chromosomal regions and genes was not drawn in scale. (Left, vertical tracks) Fifteen tumor samples were grouped as having *CCND1* amplification, and ten were confirmed to be driven by ecDNA. (Right, vertical track) *RB1* mutation indicates the predicted functional impact of detected *RB1* variants per the Ensembl Variant Effect Predictor (VEP). **(c)** ecDNA events (up to 3MB) in post-chemotherapy (post-chemo) tumors had a significantly higher mean junction copy number (JCN) and maximum JCN than ecDNA events in pre-chemotherapy (pre-chemo) tumors, indicating the increasing ecDNA complexity following systemic chemotherapy. Two-sided Wilcoxon rank-sum test. *: *P* < 0.05. Boxplots show the median and IQR. The lower whisker indicates Q1-1.5*IQR. The upper whisker indicates Q3+1.5*IQR. Each dot represents one ecDNA event. **(d – g)** Combined genome graphs of AmpliconArchitect (top track) with JaBbA (CN and read depth tracks). AmpliconArchitect classifies whether an amplicon is cyclic or non-cyclic. JaBbA tracks show the chromosomal locations and copy-number alteration for DNA segments (black vertices) and the corresponding JaBbA SV events (colored edges) in WCMIV091 (**d, e**) and WCMIV076 (**f, g**) before and after systemic therapy. WCMIV091 was treated with neoadjuvant abemaciclib, a CDK4/6 inhibitor, for four weeks as part of a clinical trial. WCMIV076 received a neoadjuvant combination of gemcitabine and cisplatin. (Top right schematics) In WCMIV091, *FGFR1* was rearranged with *CCND1* on the same ecDNA, while in WCMIV076, *FGF19* and *CCND1* were co-amplified together on ecDNA. The chromosomal locations of *CCND1, FGFR1, and FGF19* are highlighted in red. CN: copy-number, Chr: Chromosome, Gem: gemcitabine, Cis: cisplatin, BFB: breakage-fusion-bridge, TIC: templated-insertion chain.

*CCND1* encodes cyclin D1, which forms a complex with CDK4/6 to activate E2F transcription factors by phosphorylating Rb (a native inhibitor of E2F), allowing cell cycle progression through the G1/S checkpoint^37, 38^. p16 is a tumor suppressor protein encoded by cyclin-dependent kinase inhibitor 2A (*CDKN2A*) that inhibits CDK4/6 **(Fig. 4b & Supplementary Table 12)**. In samples harboring *CCND1* ecDNA amplification, 87% *CDKN2A* was deleted (mean CN fold-change relative to ploidy 0.34) **(Fig. 4b & Supplementary Table 12),** whereas downstream signaling genes *CDK4*, *CDK6*, and *RB1* were predominantly wild-type. Our results suggest that the p16-cyclinD1-CDK4/6-Rb pathway is hyperactivated in UC tumors with ecDNA *CCND1* amplification.

To gain insight into the role of ecDNA in treatment resistance, we then investigated the dynamics of ecDNA events in matched tumors undergoing systemic therapy. For ecDNA events up to 3MB in size, post-chemotherapy tumors had a significantly higher mean JCN (*P* = 0.016, Two-sided Wilcoxon rank-sum test) and max JCN (*P* = 0.032, Two-sided Wilcoxon rank-sum test) than that of chemotherapy-naïve tumors, indicating the overall increasing genetic complexity of ecDNA following treatments **(Fig. 4c)**. Specifically, *CCND1* ecDNA events increased their integer CN reflecting the mean CN of ecDNAs per cell from 17 to 26 in patient WCMIV091 and from 12 to 24 in patient WCMIV076 following systemic treatment **(Fig. 4d-g)**. These increases are consistent with mathematical models predicting ecDNA behavior under positive selection^28^. AmpliconArchitect reconstruction of ecDNA structures showed co-amplification of *FGFR1* and *FGF19* with *CCND1* in WCMIV091 and WCMIV076, respectively, and confirmed their copy number increases following systemic therapy **(Fig. 4d-g)**. Collectively, these data show the high prevalence of ecDNA-mediated *CCND1* amplification in UC and highlight the role of these events as drivers of drug resistance.

## Discussion

In this study, we employed whole-genome sequencing in advanced UC over each patient’s lifespan using matched patient samples collected from distinct anatomical regions and consecutive time points. In doing so, we unveiled a previously uncharted mutational timeline and chronology of two key mutagenic factors, APOBEC3 cytidine deaminases, and platinum-based chemotherapy. Our study provides a comprehensive map of complex SVs in advanced UC, showing a high prevalence of ecDNA amplification during UC progression. We found that ecDNA was mutagenized by APOBEC3 and chemotherapy. Furthermore, we discovered that *CCND1* ecDNA amplicons undergo dynamic evolution, incorporating additional DNA segments^29, 31, 39^, serve as adaptive reservoirs for oncogene amplification^40^, and potentially contribute to resistance against systemic therapies.

Previous pan-cancer studies have evaluated the timing and clonality of mutagenic processes of primary tumors^4, 15, 41^. However, due to the paucity of longitudinal samples, understanding these processes’ trajectories still needs to be completed. By superimposing mutational signatures on phylogenetic trees to reconstruct the tumor trajectory based on analyzing multiple tumors from the same patient, we examined the timing and clonality of APOBEC3 and chemotherapy-induced mutagenic processes in serial samples from each patient using metastatic biopsies and rapid autopsies. Our study demonstrates that APOBEC3-induced mutagenesis occurs early in the natural history of UC. The relative timing and contributions of different mutagenic processes are cancer-type specific^15^. Our results are aligned with the relatively early timing of SBS2/13 mutations in UC compared to other tumor types in the PCAWG and TCGA datasets^15, 22, 42, 43^. Furthermore, previous studies show that APOBEC3-induced mutations are found in histologically normal bladder urothelium. Our findings show a high proportion of APOBEC3-induced nonsynonymous mutations with a dN/dS ratio > 10 in key oncogenes and tumor suppressor genes, indicating clonal selection over a long period, probably starting since normal or pre-cancerous stages. This suggests these mutations confer a fitness benefit for outgrowing normal wild-type urothelial cells, ultimately driving carcinogenesis. These data are consistent with our recently published work showing that the transgenic expression of human APOBEC3G promotes mutagenesis, genomic instability, and kataegis, leading to shorter survival in a murine bladder cancer model^44^. Additional studies have demonstrated that the transgenic expression of APOBEC3 enzymes drives tumor formation *in vivo* in colon and lung cancers^45, 46^ and metastatic spread in pancreatic cancer^47^. Together, these findings highlight the emerging role of mutagenesis induced by the APOBEC3 enzymes as putative drivers of tumorigenesis. Our data open unexploited opportunities for cancer control through early detection of actionable APOBEC3-induced driver mutations in the normal urothelium in patients at a higher risk of developing UC and for intercepting this process at the pre-cancerous stages.

We and others previously described chemotherapy-induced mutations using whole-exome sequencing^9, 48^. In the current study, we quantified the rate of chemotherapy-induced mutations comprising single base and doublet base substitutions. Strikingly, we identified mutational bursts generating more than 700 mutations per month of exposure. These data are aligned with a previous study showing the accumulation of hundreds of mutations per month with up to 100 times more mutations than the aging signatures during the same period of exposure in metastatic tumors treated with platinum-based chemotherapy^1^. These findings add to our understanding of the often-underappreciated role of DNA-damaging chemotherapy as a significant mutagenic force that shapes cancer evolution.

By studying the interactions between different facets of cancer genomic instability, including mutagenesis and higher-order structural variants, we gained novel insights into the temporal order and the potential underlying mechanisms of these phenomena. We identified a significant burden of complex SV in our UC cohort. We focused on ecDNA, circular fragments of DNA that can exist independently of the chromosomes, drive tumor evolution, and introduce an additional layer of cancer heterogeneity, posing therapy resistance challenges^28, 36, 49, 50^. Using JaBbA^10^ and AmpliconArchitect^32^, we identified a high prevalence of the tyfonas^10, 33^, BFBs^24, 26, 51–53^, and DMs^24, 54, 55^ complex SVs as a source of ecDNA events. The mechanisms of ecDNA biogenesis are not fully understood^56, 57^, but the occurrence of double-stranded DNA breaks followed by a variety of repair mechanisms by homologous recombination (HR)^58^, non-homologous end joining (NHEJ)^59, 60^ or microhomology-mediated end joining (MMEJ)^61^ are thought to be critical for ecDNA formation. APOBEC3-induced deamination of genomic cytosines is known to induce double-stranded DNA breaks^7, 62^, and our findings of topographical overlap between APOBEC3-induced mutations and ecDNA junctions, as well as the early relative timing of APOBEC3-induced mutagenesis in ecDNA compared to aging and chemotherapy, suggest a potential role for APOBEC3-induced genomic instability in ecDNA biogenesis. To our knowledge, this is the first report showing platinum chemotherapy-induced mutations in ecDNA structures. This novel finding suggests that the interaction between ecDNA and chemotherapy-induced mutagenesis can compound subclonal diversification and cancer heterogeneity. As ecDNA is asymmetrically inherited in cancer cells^28, 29^, bulk WGS offers only a limited resolution for tracking complex SV and ecDNA distribution at a single-cell resolution. Emerging techniques such as single-cell Circle-seq^63^ can potentially characterize the distribution of ecDNA mutagenized by APOBEC3 and chemotherapy in tumor subclones as drivers of aggressive cancer phenotypes arising from cancer evolution.

In our UC cohort, we discovered that ecDNA formation is the most common mechanism for generating *CCND1* copy number amplification. We confirmed that UC has the highest *CCND1* ecDNA events among tumor types in the TCGA/PCAWG pan-cancer cohort. These data suggest a prominent role of ecDNA formation in driving *CCND1* copy number amplification events which are common in UC^64–66^. As *CCND1* ecDNA amplification events were found to co-occur with *CDKN2A*^del^ *RB1*^WT^ tumors, we predict that *CCND1*^ecDNA-amp^ *CDKN2A*^del^ *RB1*^WT^ genotypic configuration results in overactivation of the downstream CDK4/6 pathway, promoting cell cycle progression. Importantly, we discovered that ecDNA amplicons containing *CCND1* evolve following systemic therapy, increasing their complexity by integrating additional DNA segments. This is consistent with emerging literature suggesting that ecDNA acts as a dynamic adaptive reservoir^40^ for oncogene amplification^29, 31, 39^. Novel strategies, including the use of inhibitors of DNA-dependent protein kinases as critical components of the NHEJ pathway, can limit ecDNA size and inhibit the evolution of resistance to targeted therapy in melanomas^39^. Similar strategies are potentially applicable for preventing ecDNA-driven resistance in UC.

The increased copy number of *CCND1* we observed in post-treatment tumors compared to treatment-naïve tumors highlights the potential role of *CCND1* ecDNA as a potential mechanism of resistance to selective therapeutic pressure from cytotoxic therapy. Furthermore, *CCND1* amplification is associated with the immunosuppressive tumor microenvironment, resulting in lower response rates to immune checkpoint inhibitors in UC tumors^64, 66^. These two observations suggest that the ecDNA *CCND1* amplification is a critical mechanism of resistance to cancer cell-directed therapy and immunotherapy. These data suggest that targeted therapies inhibiting the CDK4/6 pathway in UC patients with *CCND1* ecDNA amplification could lead to significant advances in treating patients with UC. One example is effectively targeting CDK4/6 in patients selected based on their *CCND1* amplification status, such as in mantle cell lymphoma^67^. Our group is currently leading the CLONEVO window-of-opportunity clinical trial (NCT03837821) of the CDK4/6 inhibitor abemaciclib in patients with UC, which will shed light on the feasibility of this approach.

In summary, our study is a pioneering investigation into the intricate interactions between endogenous and therapy-induced mutagenic mechanisms and ecDNA in the context of UC evolution. These novel insights hold promise for future research and the development of innovative therapeutic strategies aimed at intercepting urothelial carcinogenesis and cancer evolution.

## Methods

### Patient enrollment and tissue acquisition

All experimental procedures were carried out in accordance with approved guidelines and were approved by the Institutional Review Boards at WCM. Patients recruited to this study signed informed consent under IRB-approved protocols: WCM/New York-Presbyterian (NYP) IRB protocols for Tumor Biobanking— 0201005295, GU tumor Biobanking—1008011210, Urothelial Cancer Sequencing— 1011011386, Comprehensive Cancer Characterization by (Genomic and Transcriptomic Profiling—1007011157, and Precision Medicine—1305013903). Fresh frozen and formalin-fixed paraffin-embedded (FFPE) tissue from biopsies, cystectomy, and nephroureterectomy specimens from HGUC patients were collected. All pathology specimens were reviewed and reported by board-certified genitourinary pathologists (J.M.Mosquera) in the department of pathology at WCM/NYP. Clinical charts were reviewed to record patient demographics, tobacco use, family history of cancer, concurrent cancer, treatment history, anatomic site, pathologic grade, and stage using the tumor, node, metastasis (TNM) system.

### Rapid autopsy procedures

The Englander Institute for Precision Medicine at Weill Cornell Medicine, New York-Presbyterian, has been established to promote personalized medicine focused on molecular diagnostics and therapeutics. Patients were given the option to be enrolled in the IRB-approved rapid autopsy program. In addition, patients’ next-of-kin provided written consent before autopsy. A systematic autopsy protocol is followed where normal and malignant fresh tissue is collected, allocating samples to be snap frozen or formalin-fixed. The goal is to maximize the amount of tissue collected for research purposes. Once the tissue harvest is complete, the autopsy proceeds in accordance with the protocol established by the WCM Autopsy Service. For our current study, tissue samples from multiple sites were procured from each patient, as detailed above. After hematoxylin and eosin (H&E) evaluation and frozen slide annotation, DNA was extracted for whole-genome sequencing (WGS).

### Whole genome library preparation and sequencing

Whole genome sequencing (WGS) libraries were prepared using the KAPA Hyper PCR+ library preparation kit following the manufacturer’s instructions. Prior to starting library preparation, FFPE samples were repaired using PreCR Repair Mix (NEB), per the manufacturer’s instructions, followed by an additional DNA quantification. For library preparation, DNA was sheared using a Covaris LE220. DNA fragments were end-repaired, adenylated, and ligated to Illumina sequencing adapters. Libraries went through two post-ligation bead clean-ups, PCR amplification, and a final post-PCR bead cleanup. Final library quality was verified using the KAPA qPCR Library Quantification Kit (Roche) and Fragment Analyzer (Agilent). Libraries were normalized, pooled, and sequenced on an Illumina NovaSeq 6000 sequencer using 2 x 150bp cycles.

### WGS data preprocessing, variant calling, and annotation

The New York Genome Center v6 somatic pipeline^1^ was used to align the data and call variants. Briefly, sequencing reads were aligned to GRCh38 with BWA-MEM (v0.7.15)^2^. Short alignments were removed with NYGC ShortAlignmentMarking (v2.1) (https://github.com/nygenome/nygc-short-alignment-marking), and mate-pair information was added with GATK FixMateInformation (v4.1.0)^3^. Individual lane BAMs were merged and sorted simultaneously with Novosort markDuplicates (v1.03.01), followed by GATK BQSR. SNVs, MNVs, and Indels were called with MuTect2 (GATK v4.0.5.1)^4^, Strelka2 (v2.9.3)^5^, and Lancet (v1.0.7)^6^. SVs were called with Svaba (v0.2.1)^7^, Manta (v1.4.0)^8^, and Lumpy (v0.2.13)^9^. Svaba was to call both Indels and SVs. Split-read support for SVs was quantified using SplazerS (v1.1)^10^. Germline variants were called on the matched normals with GATK HaplotypeCaller (v3.5) and filtered with GATK VQSR at tranche 99.6%. The positions of heterozygous germline variants were used to compute B-allele frequencies in the tumor samples. Variants were merged across callers and annotated with Ensembl (v93)^11^, COSMIC (v86)^12^, 1000Genomes (Phase3)^13^, ClinVar (201706)^14^, PolyPhen (v2.2.2)^15^, SIFT (v5.2.2)^16^, FATHMM (v2.1)^17^, gnomAD (r2.0.1)^18^ and dbSNP (v150)^19^ using Variant Effect Predictor (v93.2)^20^. Somatic variants that occurred in two or more individuals in our in-house Panel of Normals (PON) were removed, as well as SNV/Indels that had MAF greater than or equal to 1% in 1000Genomes or gnomAD, and SVs overlapping with DGV^21^, 1000Genomes, or gnomAD-SV^22^. SNV/INDELs with tumor VAF less than 0.0001, normal VAF greater than 0.2, depth less than 2 in either the tumor or normal, and normal VAF greater than tumor VAF were filtered from the final callset. SNV/Indels with support from 2 or more callers were marked as high confidence. SVs with support from two or more callers or one caller with split-read or CNV changepoint support were marked as high confidence. Variants detected by the somatic calling pipeline in the three FFPE normal tissues were removed from the tumor samples.

### Purity and ploidy estimation

Purity and ploidy were estimated for each sample using AscatNGS (v4.2.1)^23^ and Sequenza (v3.0.0)^24^. Estimates were manually reviewed and chosen based on VAF, BAF, and read depth fit. 6 tumors were excluded from downstream JaBbA analysis due to low purity for JaBbA purposes.

### Study sample size definition

Our study includes 79 samples. 77 histologically-proven urothelial cancers and two morphologically normal urothelial samples from 50 patients. Two normal urothelium samples from WCMIV001 and WCMIV013 were excluded from all analyses except as specific cases with reliable ploidy and purity to study the longitudinal analyses of APOBEC3 mutagenesis in the normal urothelium.

### Union of somatic variants for patients with multiple samples

For patients with multiple samples, a union of ‘HighConfidence’ somatic SNVs and Indels across all the patient’s samples was generated. Pileup (0.15.0) (https://github.com/pysam-developers/pysam) was then run on tumor and normal bam files to compute the counts for variants present in the union vcf that were missing from each sample’s vcf. Variants that had a VAF>0 were then rescued and added to the sample vcf. The resulting ‘union vcf’ was used for further post-processing.

### Mutational signature fitting and assignment

Mutational signature fitting was performed using the deconstructSigs R package (1.9)^25^. DeconstructSigs was run with ‘HighConfidence’ variants as input and COSMIC v3.2 signatures (https://cancer.sanger.ac.uk/signatures/)^12^ as a reference. The tool was run with the arguments “contexts.needed=TRUE” and “signature.cutoff=0” for SBS, DBS, and ID signatures. Following signature fitting, for each SNV, we computed a channel of 78 posterior probabilities (corresponding to each of the SBS reference signatures) that the mutation was caused by a given COSMIC signature (similar to *Kasar et al. 2015*^26^) **(equation).** Posterior probabilities less than 0.5 were discarded, and the most likely signature was chosen. If the trinucleotide context was not one of the top 5 peaks in the reference signature, the assignment was discarded. APOBEC (SBS2 and SBS13) and platinum chemotherapy (SBS31 and SBS35) were treated separately based on prior knowledge (SBS2: C>T; SBS13: C>A, C>G; SBS31: C>T, T>A; SBS35: C>A, C>G, C>T, T>A). We then leveraged patient clinical data to further refine platinum chemotherapy assignments. If a sample was taken from a patient that did not receive platinum chemotherapy, any SBS31 and SBS35 mutation assignments were removed. For patients with multiple samples, SBS31 and SBS35 mutation assignments were removed for mutations occurring in both pre-and post-chemotherapy samples. However, if a mutation was unassigned in one post-chemotherapy sample but was assigned to SBS31 or SBS35 in another post-chemotherapy sample from the same patient, the mutation in the first sample was reassigned.

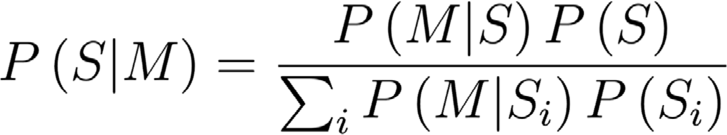

### Mutational signature assignment in matched urothelium samples

Mutational signature assignments of variants in the normal urothelium were determined using matched tumors from the same patient. Pileup (0.15.0) (https://github.com/pysam-developers/pysam) was run on the normal urothelium, using positions with an SBS2 or SBS13 assignment in the matched tumors. Variants with VAF greater than 0 were counted towards the number of variants assigned to a given mutational signature in the normal urothelium.

### Signature clonality fold change analysis

To investigate the timing of aging (SBS1), APOBEC (SBS2/13), and chemotherapy (SBS31/35) mutational processes, we computed a signature clonality fold change as described in *Pich et al. 2019*^27^. To compute the clonal vs. subclonal fold change for each sample, we pooled all MutationTimer^28^ clonal categories (early clonal, late clonal, clonal [NA], subclonal) and divided the proportion of clonal mutations assigned to the signature of interest by the proportion of subclonal mutations assigned to the same signature. The early vs. late fold change was calculated in a similar manner. Mutation counts were pooled for their respective COSMIC signatures.

### Estimating the velocity of mutagenic processes

Mutagenic velocity (rate of signature accumulation per month of exposure) was calculated for aging (SBS1), APOBEC (SBS2/13, DBS11), and platinum chemotherapy (SBS31/35, DBS5) signatures. Mutation counts from signatures fit using deconstructSigs^25^ were divided by the exposure time to mutagenic processes (in months). The estimated exposure time for aging and chemotherapy was between the date of sample collection and the date of birth, and the date of the initial chemotherapy treatment, respectively. For APOBEC, the exposure time is assumed to be from the sample collection date dated back to (1) 10 years before the UC diagnosis date or (2) the date of birth, as we expected that APOBEC mutagenesis occurs long before the diagnosis of urothelial cancer. Samples with zero APOBEC signatures contribution and post-chemotherapy samples with zero chemotherapy contribution were excluded from the calculation.

### Detection of complex structural variants with JaBbA

Read counts were corrected for GC% and mappability in 1KB bins using fragCounter (https://github.com/mskilab/fragCounter) for all tumor and normal samples. A coverage panel of normal (PON) was built from the normal samples and used to denoise the tumor coverage data using dryclean^29^. Denoised tumor coverage profiles, B-allele frequencies, and high-confidence SVs were used as input to JaBbA^30^, an algorithm that integrates CNV and SVs into a junction-balanced genome graph, computing integer copy number for both. Default parameters were used, except for the slack penalty, which was increased to 1000. Simple inversions, translocations, duplications, and deletions, as well as templated insertion chains, quasi-reciprocal pairs, rigma, pyrgo, tyfonas, breakage-fusion-bridge cycles, and double minutes, were called on the junction-balanced genome graph using the JaBbA companion R package gGnome^30^. Using the integer copy number as output by JaBbA, we computed the fraction of genome altered, defined here as the proportion of autosomes not in a neutral copy state, as defined by sample ploidy. For samples with an intermediate average ploidy (fractional value between .4 and .6, e.g., 3.5), the copy-neutral state was set as the closest two integer values (e.g., for a ploidy of 3.5, the copy-neutral states would be 3 and 4). Otherwise, the copy-neutral state was set as the rounded ploidy.

### Mutation timing and CCF calculation

MutationTimeR R package (v1.00.2)^28^ was run using the union of somatic variants along with allele-specific copy number output from JaBbA, patient gender information, and previously estimated purity values. Parameter n.boot was set to 200. MutationTimer infers a multiplicity for each mutation and assigns a timing based on the multiplicity and the allele-specific copy number configuration at that locus. Using MutationTimer multiplicities, the cancer cell fraction was computed as follows^31^:

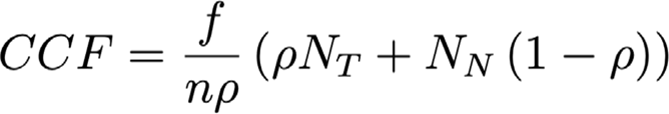

Where n is the mutation multiplicity, p is the tumor purity, f is the mutation VAF, N_T_ is the tumor total copy number at the mutation locus, and N_N_ is the normal total copy number at the mutation locus.

### Mutational signature analysis of phylogenetic trees

Phylogenetic trees were generated with the tool LICHeE (v1.0)^32^ for each patient with multiple samples. LICHeE was run with a variant by CCF matrix as input, “maxVAFAbsent” argument of 0.0, “minVAFPresent” argument of 0.5, “maxClusterDist” of 0.2, and in cell prevalence mode (-cp). The top-scoring tree was selected. The variants in each of the resulting nodes of the phylogenetic tree output were then fed through deconstructSigs as described above to estimate a set of mutational signature proportions for each node of the tree.

### Calculating the ratio of non-synonymous to synonymous substitutions (dN/dS) analysis

Genes with evidence of positive selection were detected using the R package dndscv^33^ using default parameters and GRCh38-specific reference data as supplied by the developers. As per developer instructions, mutations shared across multiple samples from the same patient were only listed once.

### Regions of recurrent structural variation

A Gamma-Poisson model, as implemented in the R package FishHook^34^, was used to discover regions of recurrent structural variation. The genome was partitioned into 100KB non-overlapping bins, and the union of breakpoints from each patient was used as input as described in *Zhou M et al. 2022*^35^. Regions overlapping intervals of mappability <1 and centromeres by more than 25% were excluded from the analysis. Covariates were added to model the background mutation rate, including:

- nucleotide frequency, dinucleotide frequency, trinucleotide frequency,
- H3K4me3 marks [ENCODE accession: ENCFF191IBA], H3K27ac marks [ENCFF208GHP], H3K4me1 marks [ENCFF759BRD], H3K3me3 marks [ENCFF983DSU],
- DNase hypersensitivity sites [ENCFF823HYK],
- replication timing (https://github.com/skandlab/MutSpot/tree/master/features/Ch38), fragile sites [HGNC 2021], and
- RepeatMasker LINE, SINE, LTR, simple repeat, and DNA transposon annotations from UCSC^36^.

An FDR-adjusted (Benjamani-Hochberg) p-value cutoff of 0.25 was used to nominate significant breakpoint hits. Cancer-related genes were from COSMIC Cancer Gene Consensus^37^.

### Detection of extrachromosomal DNA

We noted that JaBbA does not impose any kind of cyclic constraint when calling high-level amplifications (e.g., double minutes, tyfonas, and breakage-fusion-bridges). To determine whether these events were circular ecDNA, AmpliconArchitect (v1.2)^38^ was run using default parameters on tumor BAMs down-sampled to 20X, considering only intervals with integer copy numbers greater than 4 (as inferred by JaBbA) and longer than 10KB. If any high-level amplifications, as detected by JaBbA, overlapped with a cyclic amplicon, as detected by AmpliconArchitect, they were nominated as ecDNA.

### Kataegis and kyklonas identification

We ran the SigProfilerClusters^39^ software with default parameters to identify kataegis loci, computing a sample-dependent inter-substitution distance of clustered mutations and requiring a kataegis event to have a consistent VAF. Kataegis events contained completely within the footprint of an ecDNA are classified as kyklonas.

### Calculating VAF of APOBEC and chemotherapy mutations on ecDNAs

As a secondary method for determining the relative contributions and timing of mutational processes acting on ecDNA, we partitioned ecDNA SNVs by VAF into three groups (VAF ≤ 0.333, 0.333 < VAF ≤ 0.667, VAF > 0.667) for each sample. We then ran deconstructSigs as previously described for each sample and VAF combination, requiring that a particular combination have at least 50 SNVs.

### Analysis of ecDNA *CCND1* amplification in TCGA/PCAWG pan-cancer cohorts

The AmpliconArchitect analyses from *Kim et al. 2020*^40^ were accessed from GitHub. Samples were from both TCGA and PCAWG datasets (based on their IDs, either SA-XXXX or TCGA-XXXX). Sample barcodes from all specimens were analyzed for amplicon intervals corresponding to the genomic location of *CCND1* (chr11:69,455,855-69,469,242). The resulting amplicon intervals were then matched to the type of amplification event (heavily rearranged, circular-breakage fusion bridge, or linear) as determined by AmpliconArchitect. Only one *CCND1* amplification event was identified in cervical cancer and none in B-cell lymphoma, glioblastoma, sarcoma, renal, and colorectal cancer.

### Statistical tests

The two-sided Wilcoxon rank sum test, two-sided t-test, or Pearson test for continuous variables was performed using R version 4.0.0 software. Unless specified otherwise, P<0.05 was considered statistically significant.

### Ethical Approval

The studies involving human participants were reviewed and approved by Weill Cornell Medicine IRB. The patients/participants provided their written informed consent to participate in this study. Written informed consent was obtained from the individual(s) for the publication of any potentially identifiable images or data included in this article.

## Data availability statement

All BAM files and associated sample information are deposited in dbGaP under accession <u>phs001087.v4.p1</u>. The TCGA/PCAWG pan-cancer human cancer data^40^ used for *CCND1* amplification analysis was obtained and modified from the Supplementary Information of https://doi.org/10.1038/s41588-020-0678-2.

## Code availability statement

The open source code used in this paper were listed: JaBbA (https://github.com/mskilab/JaBbA), gGnome (https://github.com/mskilab/gGnome), AmpliconArchitect (https://github.com/virajbdeshpande/AmpliconArchitect), FishHook (https://github.com/mskilab/fishHook), MutationTimeR (https://github.com/gerstung-lab/MutationTimeR), deconstructSigs (https://github.com/raerose01/deconstructSigs), SigProfilerClusters (https://github.com/AlexandrovLab/SigProfilerClusters), Pileup (https://github.com/pysam-developers/pysam), ShortAlignmentMarking (https://github.com/nygenome/nygc-short-alignment-marking), BWA MEM (https://github.com/lh3/bwa), GATK (https://github.com/broadinstitute/gatk), MuTect2 (https://github.com/broadinstitute/gatk), Strelka2 (https://github.com/Illumina/strelka), Lancet (https://github.com/nygenome/lancet), Svaba (https://github.com/walaj/svaba), Manta (https://github.com/Illumina/manta), Lumpy (https://github.com/arq5x/lumpy-sv), SplazerS (https://github.com/seqan/seqan/tree/master/apps/splazers), Ensembl (https://www.ensembl.org), COSMIC (https://cancer.sanger.ac.uk), COSMIC Cancer Gene Consensus (https://cancer.sanger.ac.uk/census), ClinVar (https://www.ncbi.nlm.nih.gov/clinvar/), PolyPhen (http://genetics.bwh.harvard.edu/pph2/index.shtml), SIFT (http://sift-dna.org/sift4g), FATHMM (http://fathmm.biocompute.org.uk), gnomAD (https://gnomad.broadinstitute.org/), gnomAD-SV (https://gnomad.broadinstitute.org/,

https://github.com/talkowski-lab/gnomad-sv-pipeline), dbSNP (https://www.ncbi.nlm.nih.gov/snp/), Variant Effect Predictor (VEP) (http://www.ensembl.org/vep), Database of Genomic Variants (DGV) (http://dgv.tcag.ca/), AscatNGS (https://github.com/cancerit/ascatNgs), Sequenza (http://www.cbs.dtu.dk/biotools/sequenza), LICHeE (https://github.com/viq854/lichee), fragCounter (https://github.com/mskilab/fragCounter), dryclean (https://github.com/mskilab/dryclean), MutSpot (https://github.com/skandlab/MutSpot/tree/master/features/Ch38), RepeatMasker (https://github.com/rmhubley/RepeatMasker). Custom analysis scripts and scripts to reproduce figures are available at: https://github.com/nygenome/UrothelialCancer_WGS_paper_figures.

## Supporting information

Structural Variants Glossary

## Acknowledgments

We would like to thank our patients and their families for participating in this study. We would like to thank Paul Yoo for the constructive review of the manuscript. Work was partially supported by the Translational Research Program at WCMC Pathology and Laboratory Medicine. This work was supported by the New York Genome Center’s Polyethnic-1000 Initiative (P-1000), the National Institute of Health NIH U01 CA260369-01, Weill-Cornell Clinical & Translational Science Center CTSC Pilot Award UL1 TR002384-06, the Department of Defense W81XWH-17-1-0539, Starr Cancer Consortium SCC I14-0047, The Leo & Anne Albert Institute for Bladder Cancer Care and Research.

## Author Contributions

Initiation and design of the study: D.D. Nguyen, W.F. Hooper, O. Elemento, N. Robine, and B.M. Faltas. Subject enrollment, sample collection and preparation, and clinical data collection: D.D. Nguyen, W.F. Hooper, Z.R. Goldstein, L. Winterkorn, M. Sigouros, J. Manohar, J. Moyer, D. Wilkes, S.T. Tagawa, D.M. Nanus, J.T. Nauseef, C.N. Sternberg, A.M. Molina, D. Scherr, G. Inghirami, J.M. Mosquera, O. Elemento, N. Robine, and B.M. Faltas. Statistical and bioinformatics analyses: D.D. Nguyen, W.F. Hooper, T.R. Chu, H. Geiger, J.M. Shelton, M. Shah, R.R. Singh, W. Liu, A. Sboner, O. Elemento, N. Robine, and B.M. Faltas. Supervision of research: O. Elemento, N. Robine, and B.M. Faltas. Writing of the first draft of the manuscript: D.D. Nguyen and B.M. Faltas. All authors contributed to the writing and editing of the revised manuscript and approved the manuscript.

## Additional information

**Supplementary information** is available for this paper.

Correspondence and requests for materials should be addressed to B.M. Faltas.

## Supplementary Figure Legends

**Supplementary Figure 1. Clinical characteristics of our study cohort**. Schematic of anatomical sites of primary and metastatic urothelial cancer samples. At each site, the numbers correspond to the number of tumors stratified by chemotherapy treatment status.

**Supplementary Figure 2. Workflow for mutational signature fitting and assignment. (a)** Workflow for mutational signature fitting by deconstructSigs and our procedure developed to assign individual mutations to COSMIC v3.2 mutational signatures as described in the **Methods** section. COSMIC: Catalogue Of Somatic Mutations In Cancer, VCF: variant cell frequency, MAP: maximum a posterior, QC: quality check. **(b)** Significant correlation between the contributions of deconstructSigs-fitted mutational signatures (x-axis) and the post-assignment contribution (y-axis) of mutations attributed to APOBEC3 (SBS2/13) and platinum-based chemotherapy (SBS31/35) validates our assignment method.

**Supplementary Figure 3. The landscape of mutational signatures induced by endogenous mutagenic processes and exogenous exposures in advanced urothelial carcinoma. (a), (b), (c)** bar graphs representing the contribution of mutational signatures induced by APOBEC3, platinum chemotherapy, and tobacco smoking (COSMIC v3.2), respectively, in 77 urothelial tumors in our study. Each panel includes associated SBSs (top barplots) and DBSs (bottom barplots) attributed to each mutagenic process.

**Supplementary Figure 4. Phylogenetic trees from individual patients depicting the relative timing of APOBEC3 and chemotherapy-induced mutagenesis in urothelial cancer evolution. (a) – (m)** Superimposed mutational signatures on nodal branching points in phylogenetic trees for patients with at least two tumor samples. Each node represents the number of SNVs (center) and is annotated with the estimated cancer cell fraction (CCF) (blue text) and the mutated genes. For each node, the SNVs were used as input for deConstructSigs to estimate the proportions of SBS, DBS, and small ID mutational signatures represented by three concentric circles from the periphery to the center, respectively. The tumor samples used for phylogenetic analysis are traced back to their corresponding anatomical sites on the schematic of each patient with annotation of their chemotherapy treatment status. The numbers on the human diagram represent the number of samples collected from the tumor sites.

**Supplementary Figure 5. Relative mutagenic velocity (average mutations per month of exposure) of platinum chemotherapy-induced and APOBEC3-induced SBS and DBS signatures.** The onset of APOBEC3-induced mutagenesis was modeled to begin at different timeframes within the patient’s lifetime. All models showed lower relative mutagenic velocity compared to chemotherapy. The x-axis represents the exposure from the estimated time of mutagenesis onset until sample collection. Samples with zero APOBEC3 signatures contribution and post-chemotherapy samples with zero chemotherapy contribution were excluded. Two-sided t-test. *: *P* < 0.05. ***: *P* < 0.001. Boxplots show the median and IQR. The lower whisker indicates Q1-1.5*IQR. The upper whisker indicates Q3+1.5*IQR. Each dot represents one tumor. UC: urothelial carcinoma.

**Supplementary Figure 6. Simple and complex structural variants in urothelial carcinoma. (a)** The proportion of simple and complex structural variant (SV) events from 71 samples from 50 patients in our UC cohort. **(b)** Tyfonas events have a significantly higher junction burden than other complex SVs. Wilcoxon rank-sum test. **: *P* < 0.01. ***: *P* < 0.001. Boxplots show the median and IQR. The lower whisker indicates Q1-1.5*IQR. The upper whisker indicates Q3+1.5*IQR. Each dot represents one event.

**Supplementary Figure 7. Enrichment of structural variants in *TP53* mutant urothelial cancers. (a)** The fraction of genome altered (FGA) is significantly higher in 42 *TP53*-mutant UC tumors compared to 28 *TP53*-wild-type (WT) tumors. **(b)** 42 tumors harboring *TP53* high and moderate-impact mutations exhibited a significant increase in the number of total junctions, deletions, duplications, chromoplexy, TIC, and BFB compared to 28 *TP53*-wild-type tumors. In both panels, the Wilcoxon rank-sum test. **: *P* < 0.01. ***: *P* < 0.001. N.S.: not significant. Boxplots show the median and IQR. The lower whisker indicates Q1-1.5*IQR. The upper whisker indicates Q3+1.5*IQR. Each dot represents one tumor. BFB: breakage-fusion-bridge. TIC: templated-insertion chains. QRP: quasi-reciprocal pairs.

**Supplementary Figure 8. Complex structural variants associated with ecDNA events in urothelial cancers.** The barplot depicts the proportion of JaBbA events that overlapped with an AmpliconArchitect’s cyclic calls. BFB: breakage-fusion-bridge. TIC: templated-insertion chains, QRP: quasi-reciprocal pairs.

**Supplementary Figure 9. APOBEC3 and platinum chemotherapy-induced mutagenesis in extrachromosomal DNA.** The graphs depict 28 ecDNA events having at least one kyklonic event in addition to the events depicted in **Fig. 2g**. The top “CN” track represents the JaBbA genome graph showing the copy-number for rearranged DNA segments (gray vertices) with SV junctions (aqua blue edges) that form the circular ecDNA events in each urothelial tumor. The middle “Kyklonas" track shows the normalized VAF of identified APOBEC3-induced kyklonas. The bottom “Non-clustered” track shows the normalized VAF of non-clustered mutations assigned to both APOBEC3-associated (SBS2/13) and platinum chemotherapy-induced (SBS31/35) mutational signatures on ecDNA. Chr: chromosome.

## Description of additional supplementary data files

**Supplementary Table 1.** Clinical characteristics of patients and samples from the WCM-UC cohort.

**Supplementary Table 2.** The mutational landscape of COSMIC SBS, DBS, and ID signatures in the WCM-UC cohort.

**Supplementary Table 3.** Superimposed mutational signatures and phylogenetic trees of UC patients with at least two tumor samples.

**Supplementary Table 4.** The mutation burden and coding mutations in the normal urothelium.

**Supplementary Table 5.** dN/dS mutations of significantly mutated genes of the WCM-UC cohort.

**Supplementary Table 6.** Composite mutations of CGC gene in the WCM-UC cohort.

**Supplementary Table 7.** The SV landscape of the WCM-UC cohort. **Supplementary Table 8A.** ecDNA events detected across the WCM-UC cohort. **Supplementary Table 8B.** Clustered and non-clustered SNVs found within ecDNA.

**Supplementary Table 9.** Regions of recurrent structural variation as identified by FishHook.

**Supplementary Table 10.** *CCND1* ecDNA amplification in the WCM-UC cohort.

**Supplementary Table 11.** *CCND1* amplification events in TCGA/PCAWG pan-cancer cohorts and WCM-UC cohort.

**Supplementary Table 12.** Modal copy number of genes involved in the p16-CyclinD1-CDK4/6-Rb pathway.

