## Supplementary material for "The Interplay between Mutagenesis and Extrachromosomal DNA Shapes Urothelial Cancer Evolution": Structural Variants Glossary

**SV Glossary**

**Structural variation**

**Junction**^1^**:** Adjacency between two genomic intervals that are not contiguous in the reference genome, also commonly referred to as a breakpoint.

**Simple deletion**^1^**:** Copy loss of a genomic interval, characterized by a single junction with supporting read pairs in the +/- orientation.

**Simple duplication**^1^**:** Copy gain of a genomic interval, characterized by a single junction with supporting read pairs in the -/+ orientation.

**Simple inversion/inverted duplication**^1^**:** Inversion of a genomic interval, characterized by two junctions with supporting read pairs in the +/+ and -/- orientations. Inverted duplications have a copy gain in the inverted segment.

**Fold-back inversion**^1^**:** +/+ and -/- orientation junctions that begin and end at very close points in the genome (in JaBbA, close == <100KB)

**Simple translocation**^1^**:** Junction between two different chromosomes, with supporting read pairs in any orientation (+/-, -/+, +/+, -/-)

**Templated insertion chain (TIC)**^1^**:** Multi-junction chains linking distant loci, resulting in copy gains of the regions involved in the event.

**Chromoplexy**^2^**:** Chains of reciprocal (balanced), or nearly-reciprocal junctions linking distant loci (in JaBbA, distant == >10MB).

**Chromothripsis**^3,4^**:** Clusters of many interleaved junctions characterized by oscillating copy number, reassembled from a single shattering event.

**Breakage-fusion-bridge cycle (BFB)**^5^**:** Structural variant in which amplification occurs over multiple cycles of cell division. Telomere loss is followed by the fusion of sister chromatids during replication, which is subsequently torn apart during division, with each daughter cell receiving a chromosome lacking a telomere. This process results in a high proportion of fold-back inversions.

**Double minute**^6^**:** A synonym for extrachromosomal DNA (ecDNA). In JaBbA, these are amplicons with a low proportion of fold-back inversions.

**Structural variation (JaBbA-specific)**

**Junction copy number**^7^**:** Analogous to the copy number of a genomic locus, the junction copy number (JCN) refers to the number of copies of a rearrangement linking to loci not contiguous on the reference. For example, a duplication-like junction resulting in the gain of a single copy would have a JCN of 1. A deletion-like junction resulting in the complete loss of a locus in a diploid genome would have a JCN of 2.

**Rigma**^7^**:** Clusters of low-JCN deletions associated with late replicating regions and fragile sites.

**Pyrgo**^7^**:** Clusters of low-JCN duplications associated with early replicating regions and super-enhancers.

**Tyfonas**^7^**:** Amplicons with a high proportion of fold-back inversions, as in breakage-fusion-bridge cycles, but with a much higher genomic mass (copy number-weighted genomic footprint) and many high-JCN junctions.

**Quasi-reciprocal pair**^7^**:** A pair of junctions with a small gap (<1MB) between them, similar to chromoplexy and TIC. (<https://github.com/mskilab/gGnome/blob/master/R/eventCallers.R#L3131>)

**Structural variation (Amplicon Architect-specific)**

**Cyclic**^6,8^**:** Amplicon Architect’s classification of ecDNA. BFB- amplicons with either:

- At least one amplified cyclic path of length >100KB.
- Greater than 12% length-weighted copy number assigned to cyclic paths.
- >50KB of cyclic paths with >5KB rearrangements and copy number > 4.5.

**Complex non-cyclic**^6,8^**:** BFB- and ecDNA- amplicons with a high fraction of length-weighted copy number assigned to paths with >5KB rearrangements.

**Linear amplification**^6,8^**:** Amplicons not classified as BFB, ecDNA, or complex non-cyclic (e.g., non-cyclic amplicons with few junctions).
