## Supplementary figures and images for "The Interplay between Mutagenesis and Extrachromosomal DNA Shapes Urothelial Cancer Evolution"

### Supplementary Figure 1

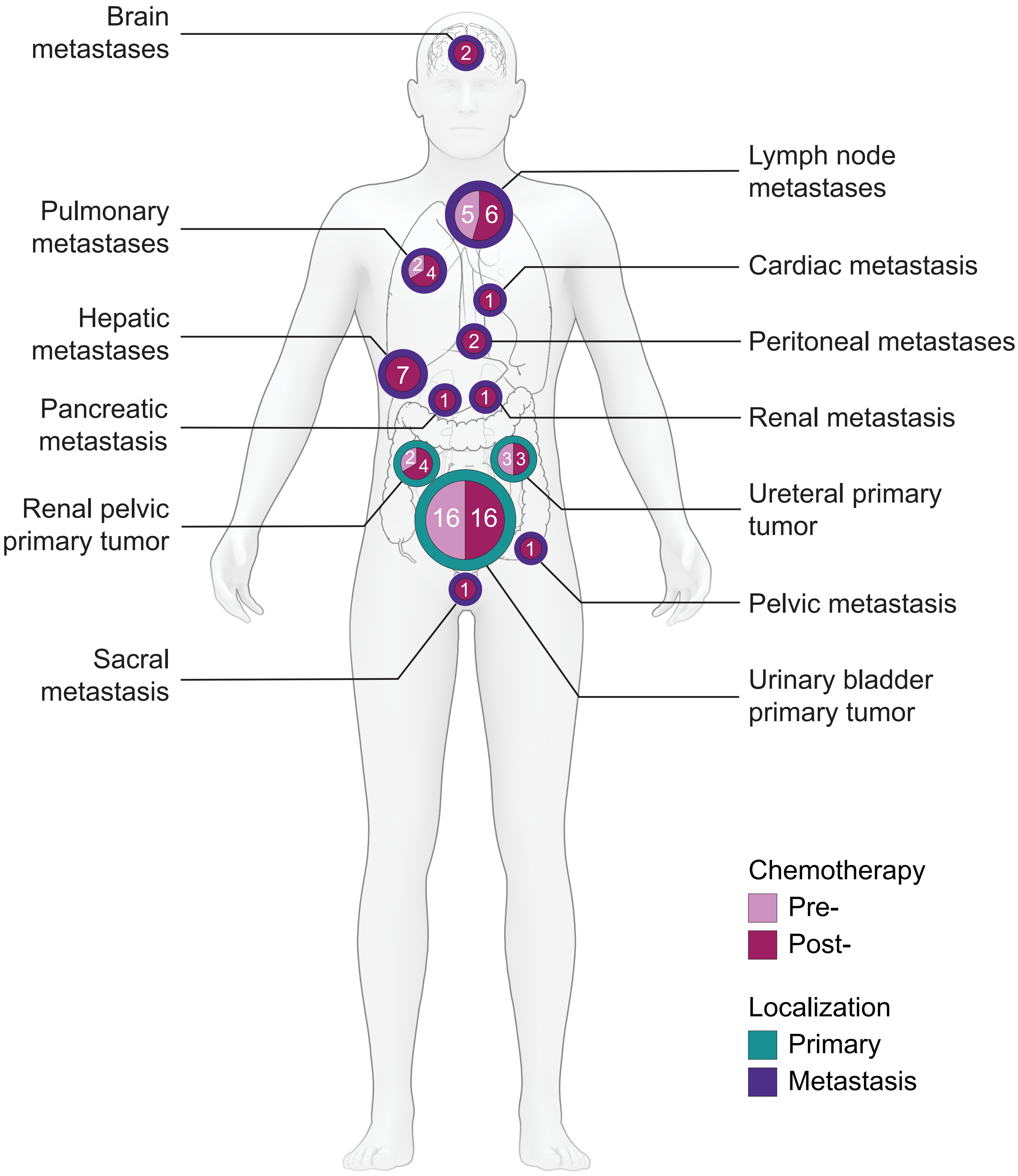

### Supplementary Figure 2

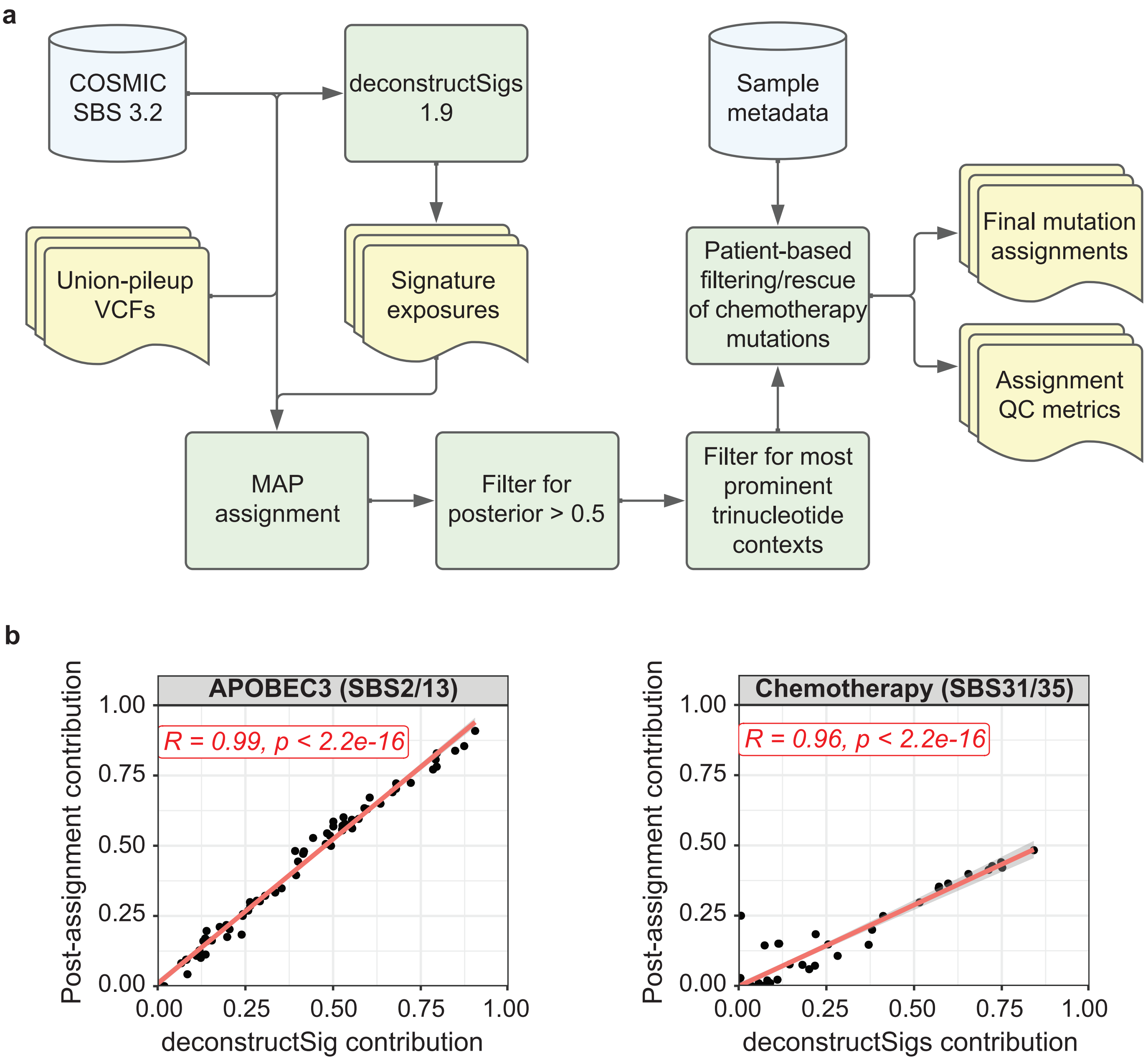

### Supplementary Figure 3

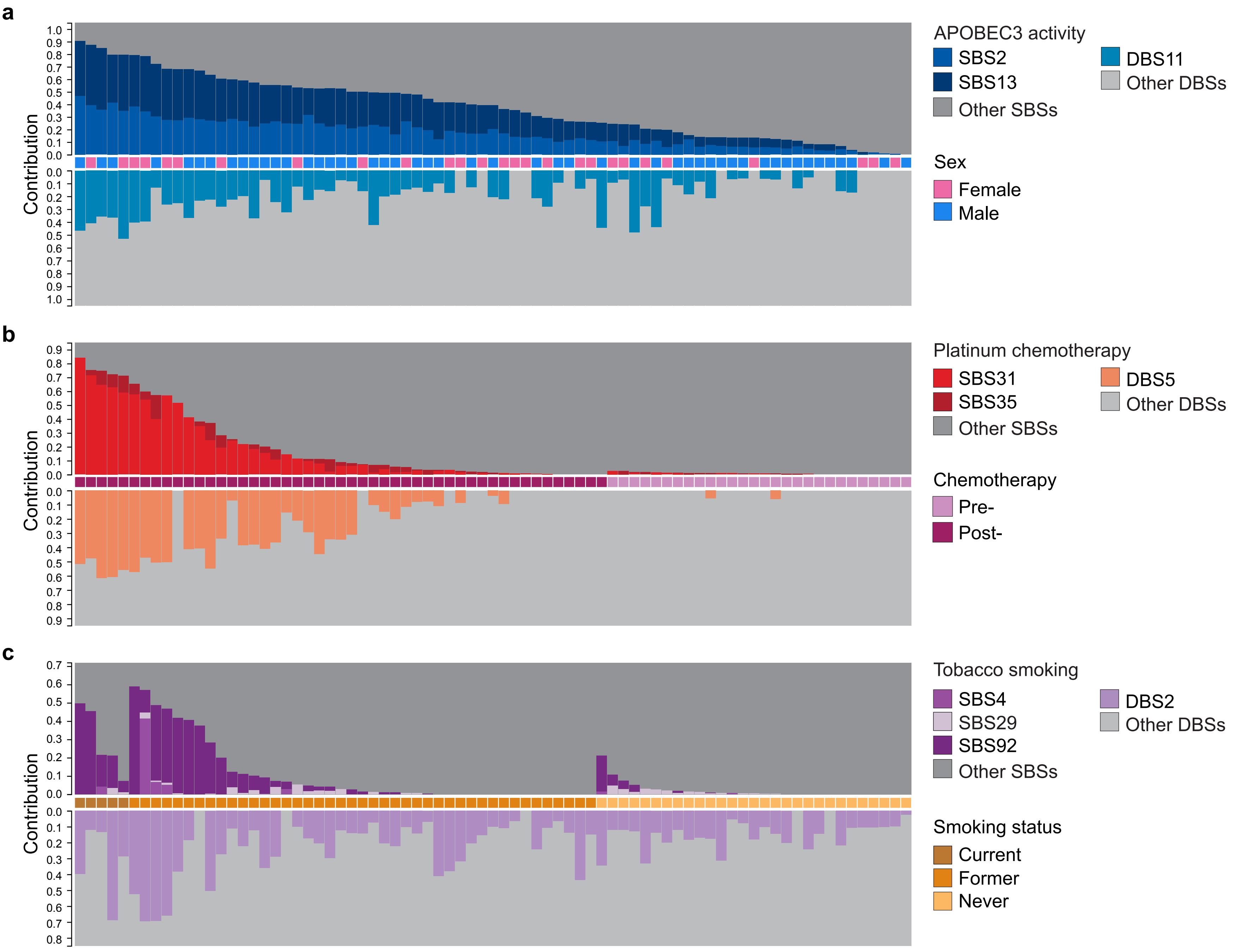

### Supplementary Figure 4a-b

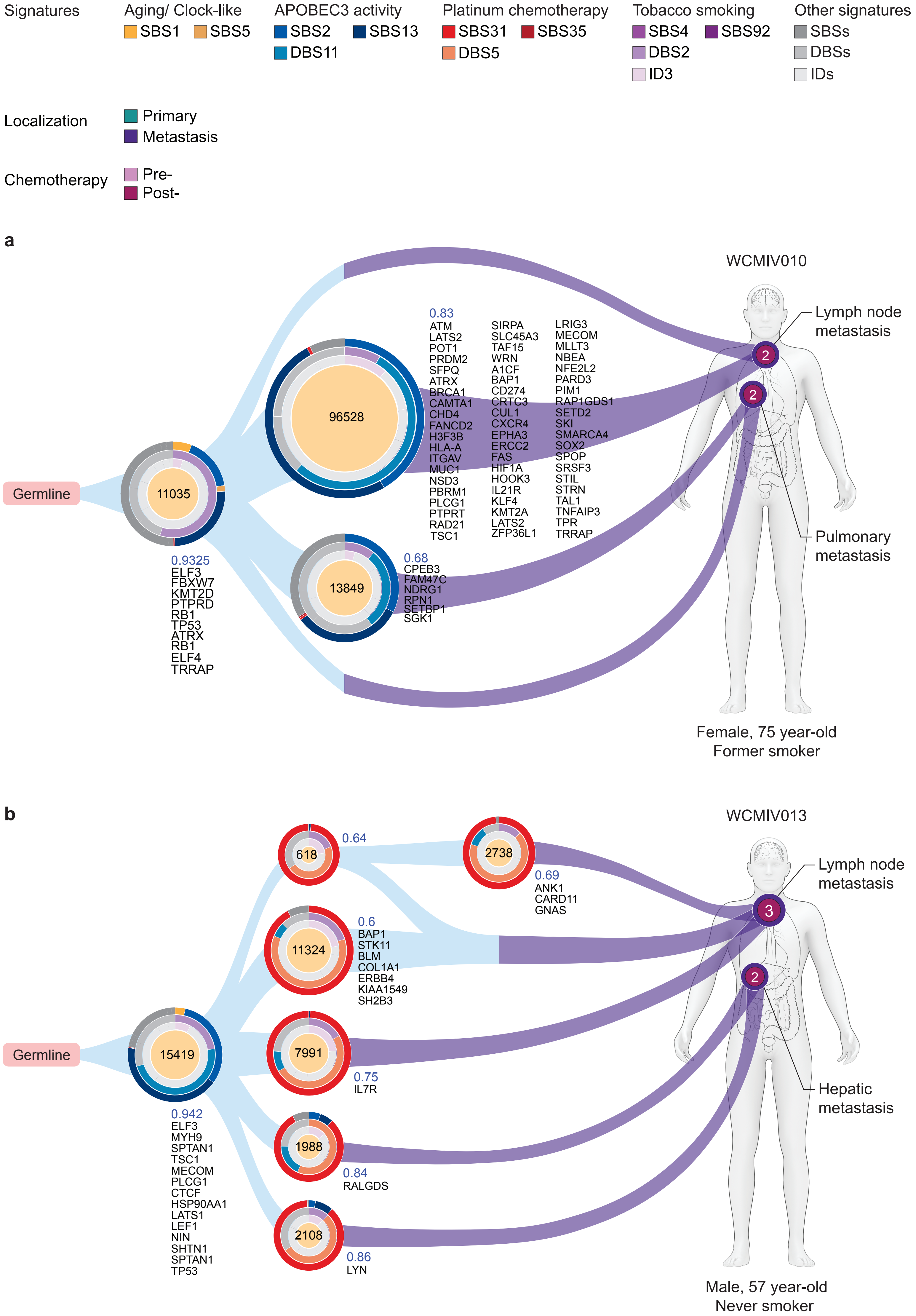

### Supplementary Figure 4c-d

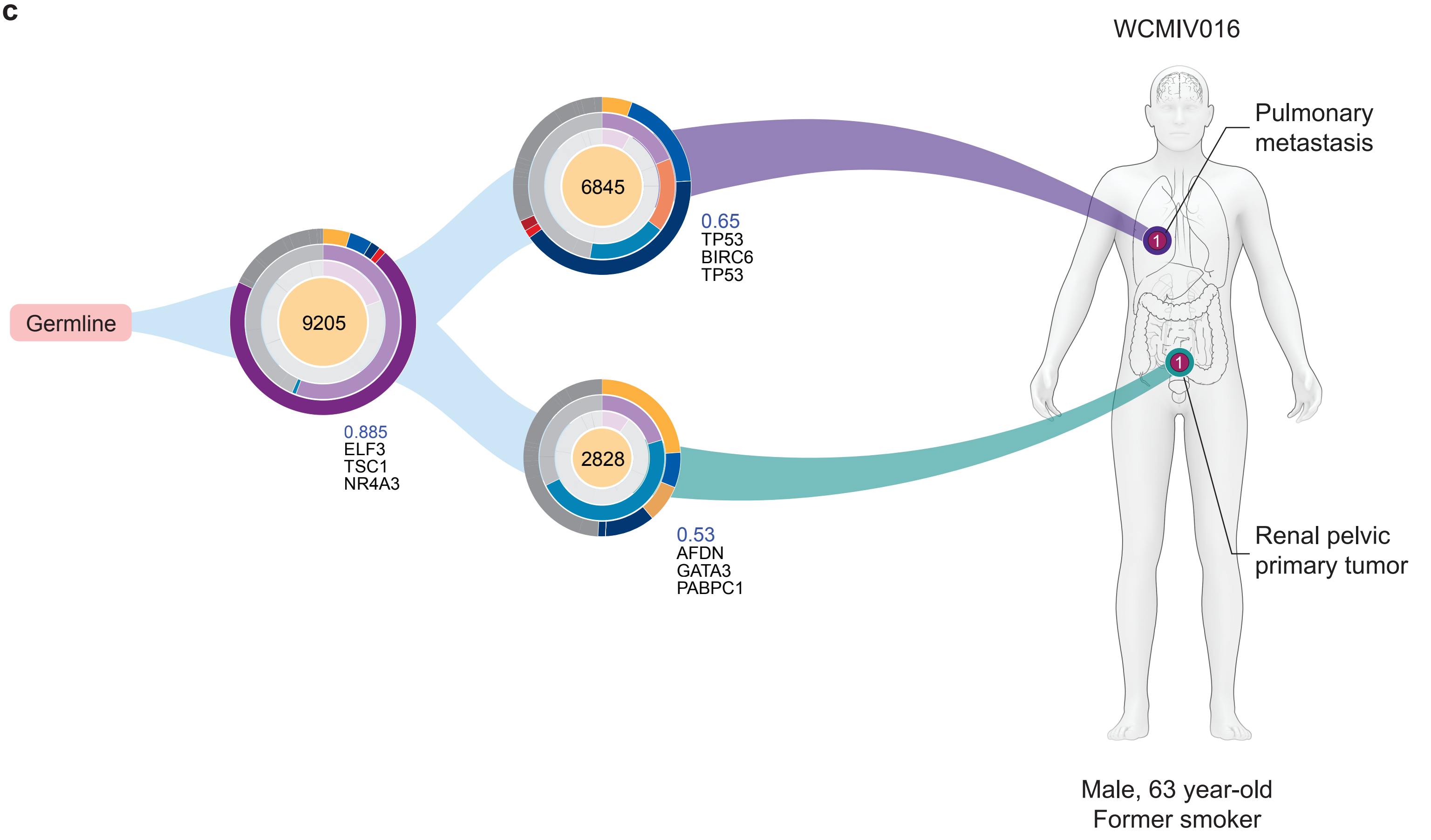

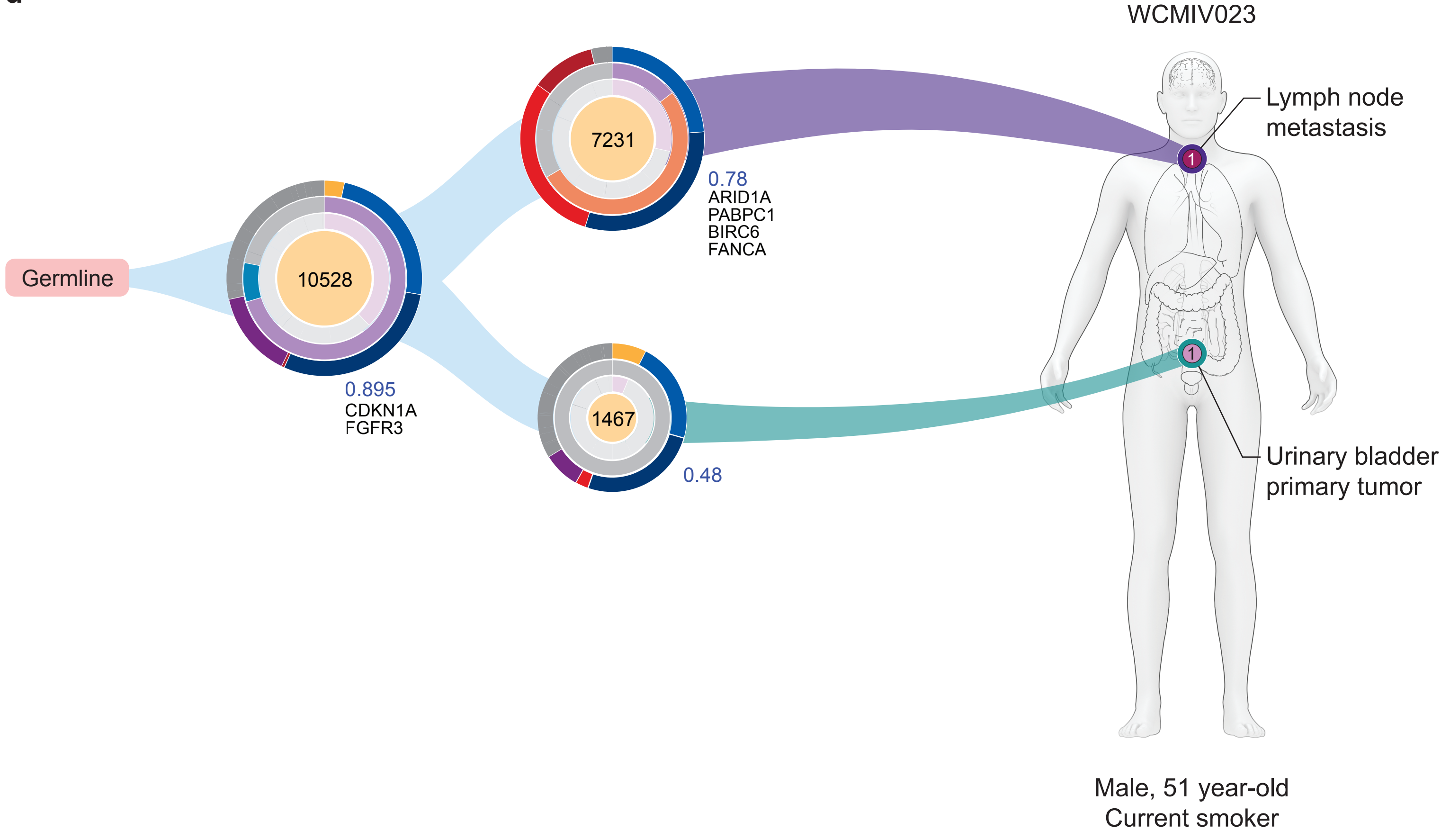

### Supplementary Figure 4e-f

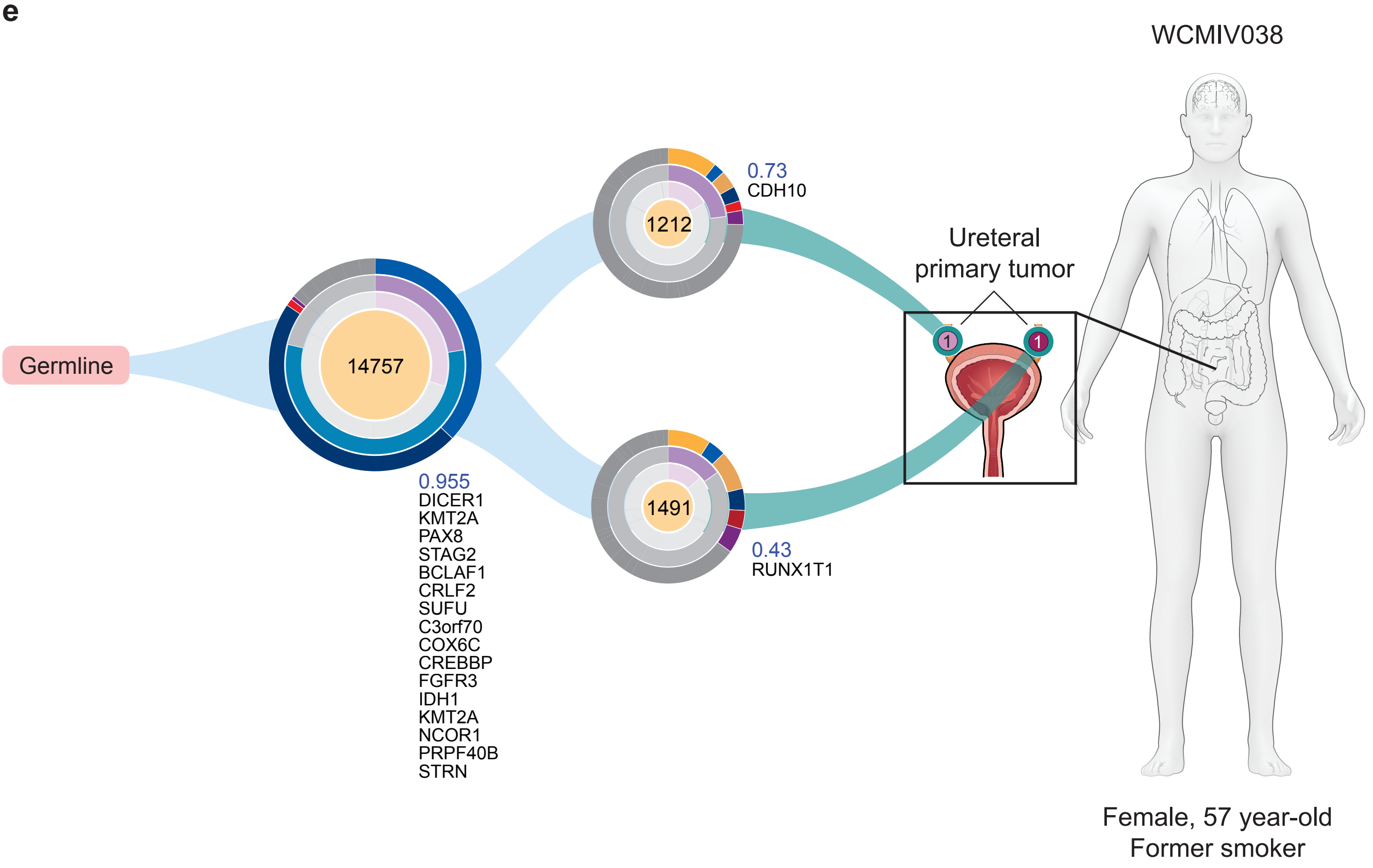

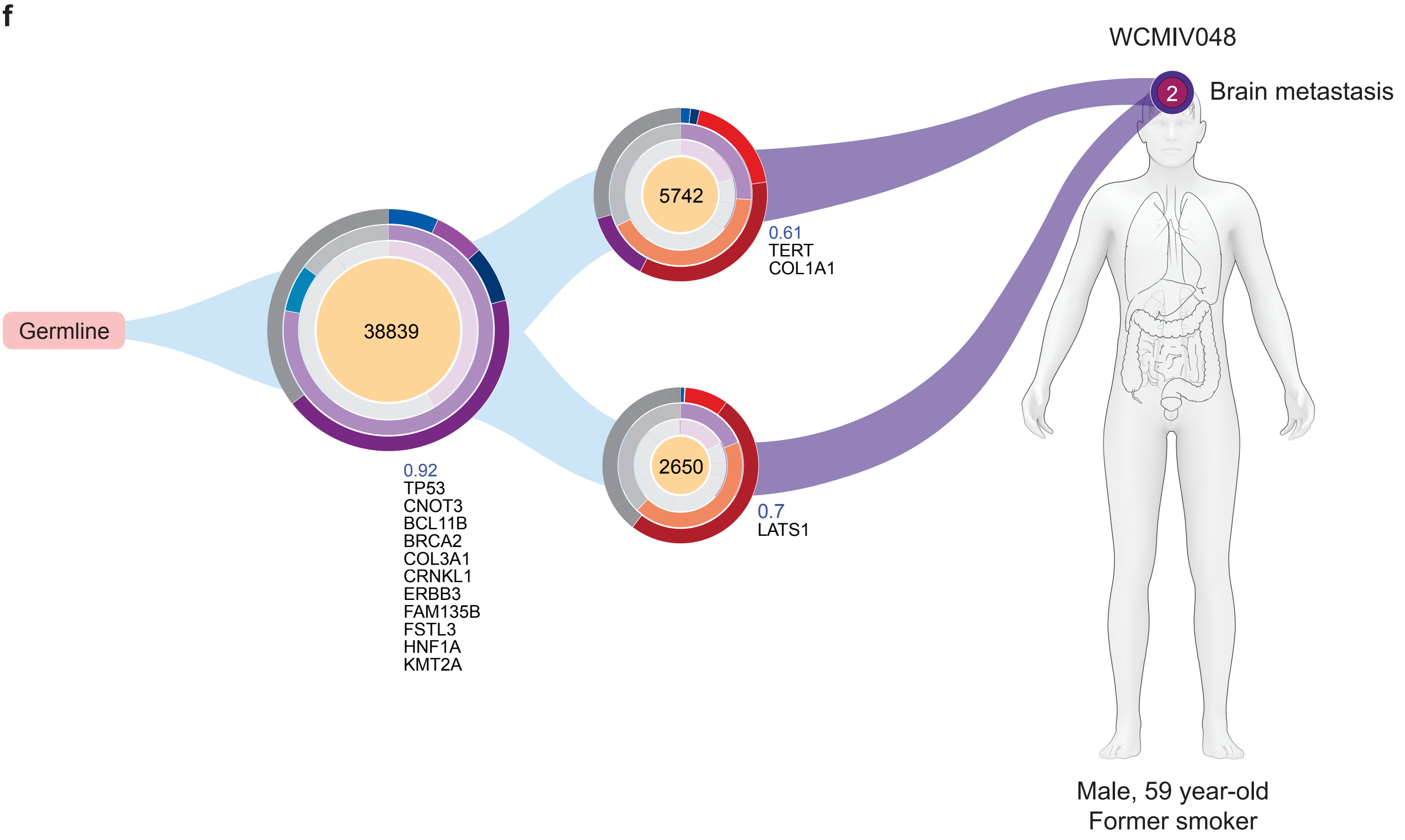

### Supplementary Figure 4g-h

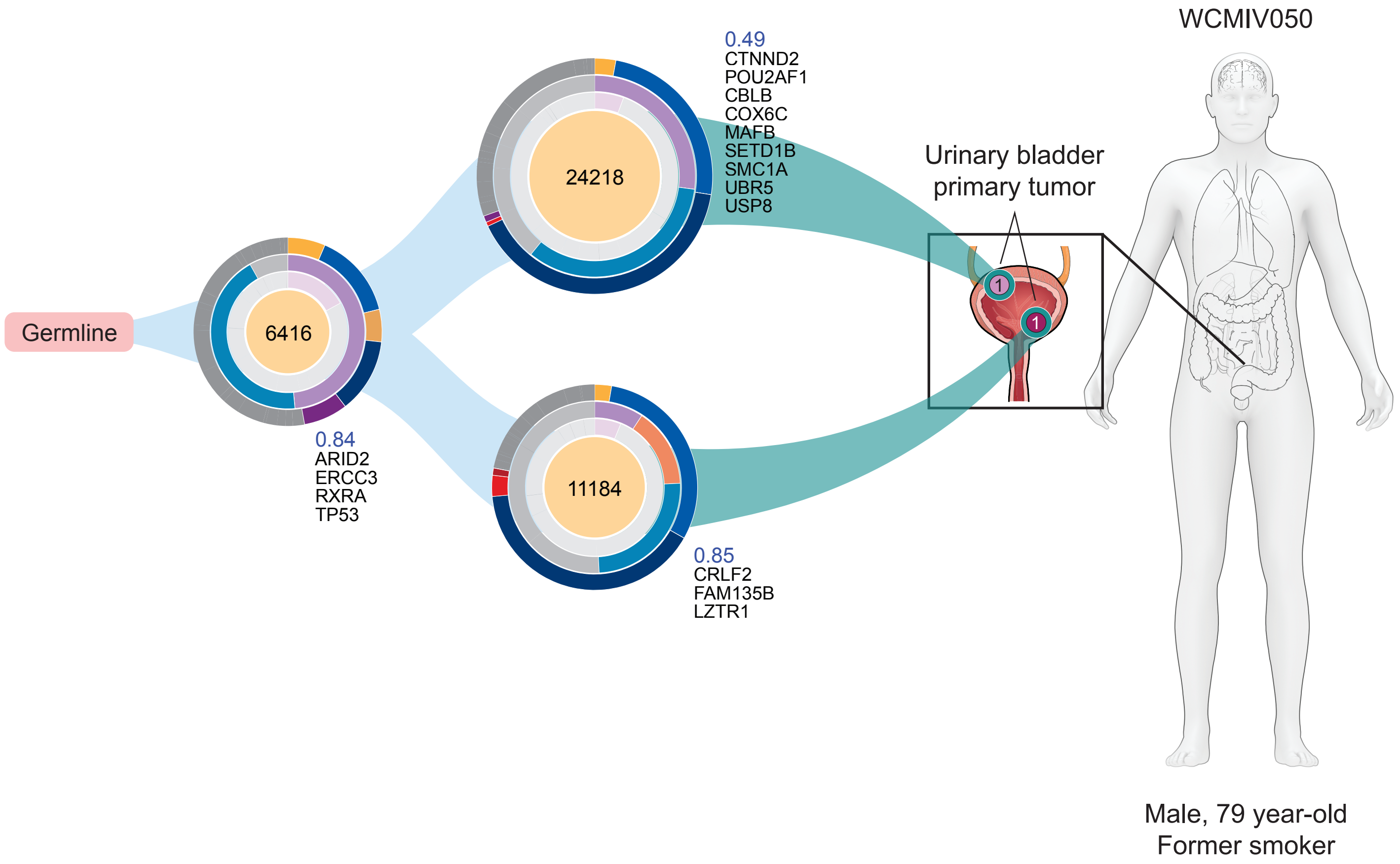

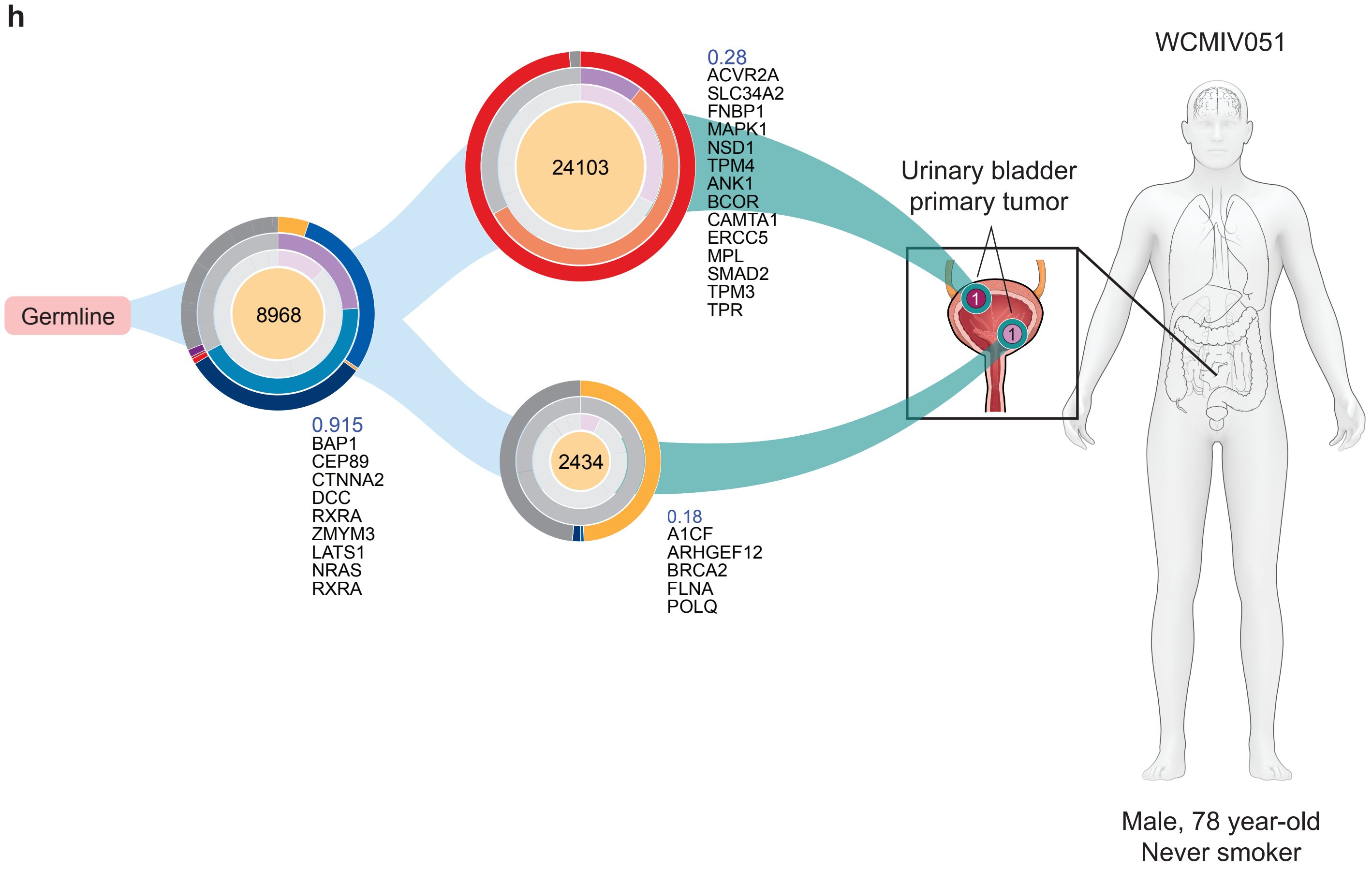

### Supplementary Figure 4i-j

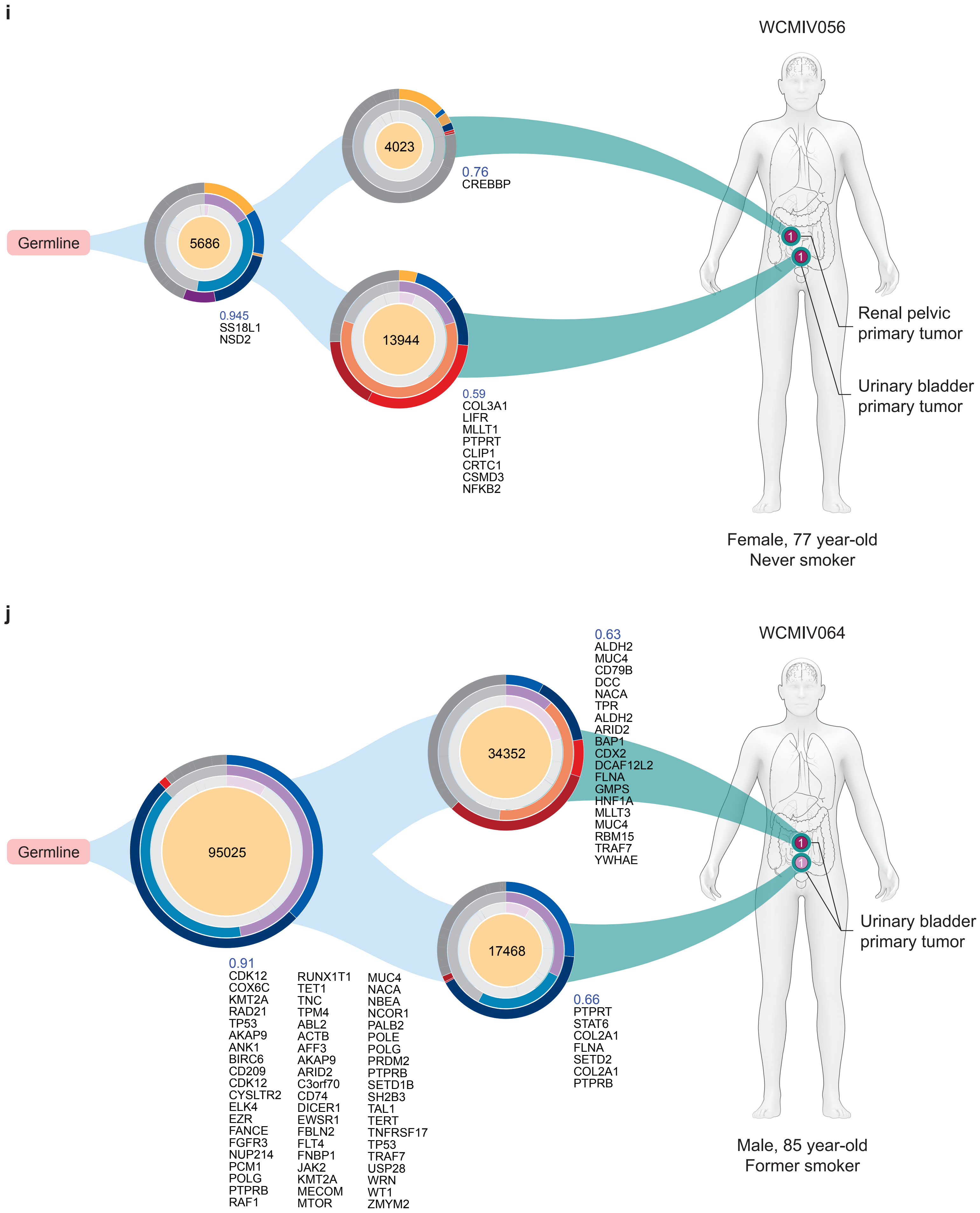

### Supplementary Figure 4k-l

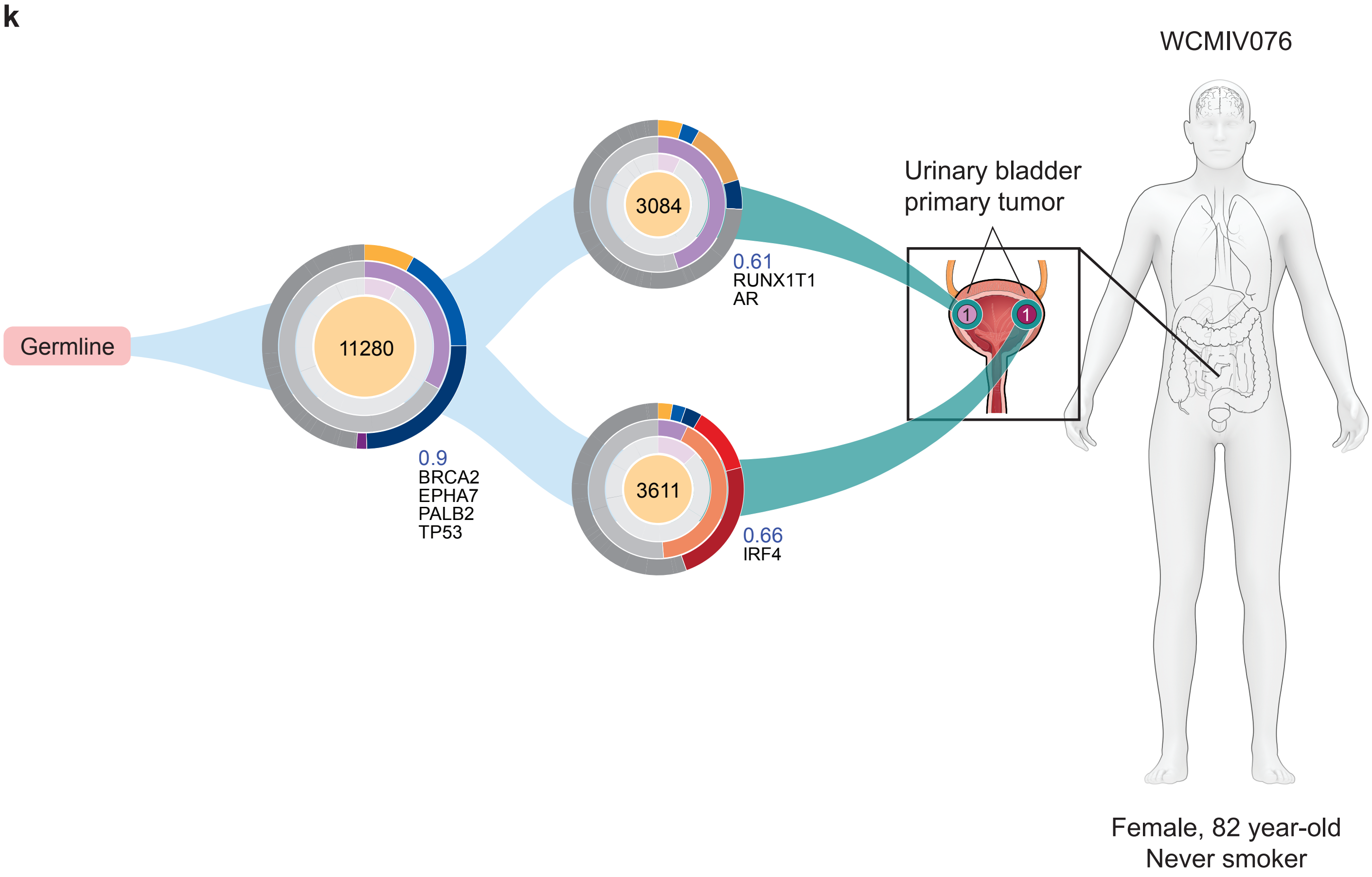

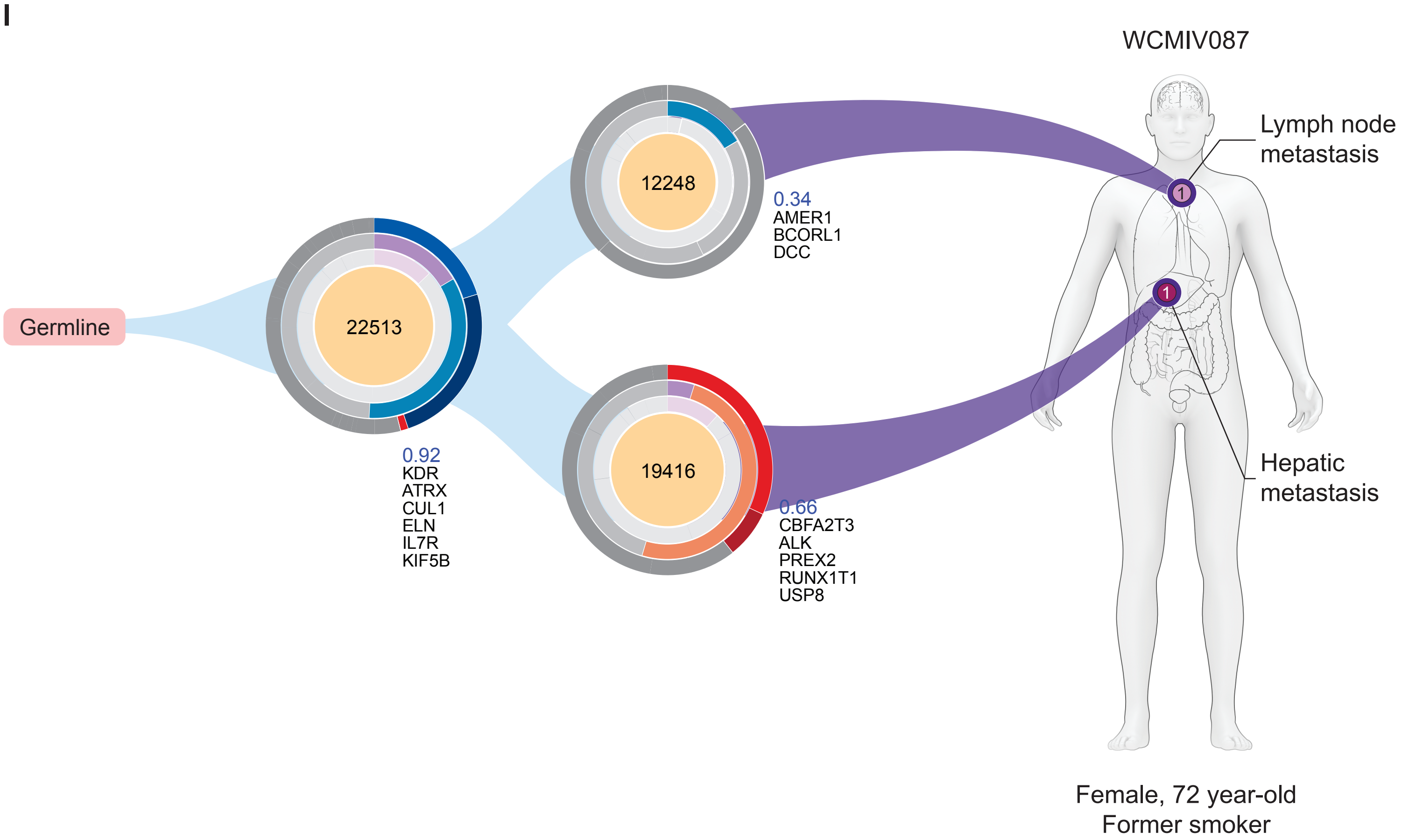

### Supplementary Figure 4m

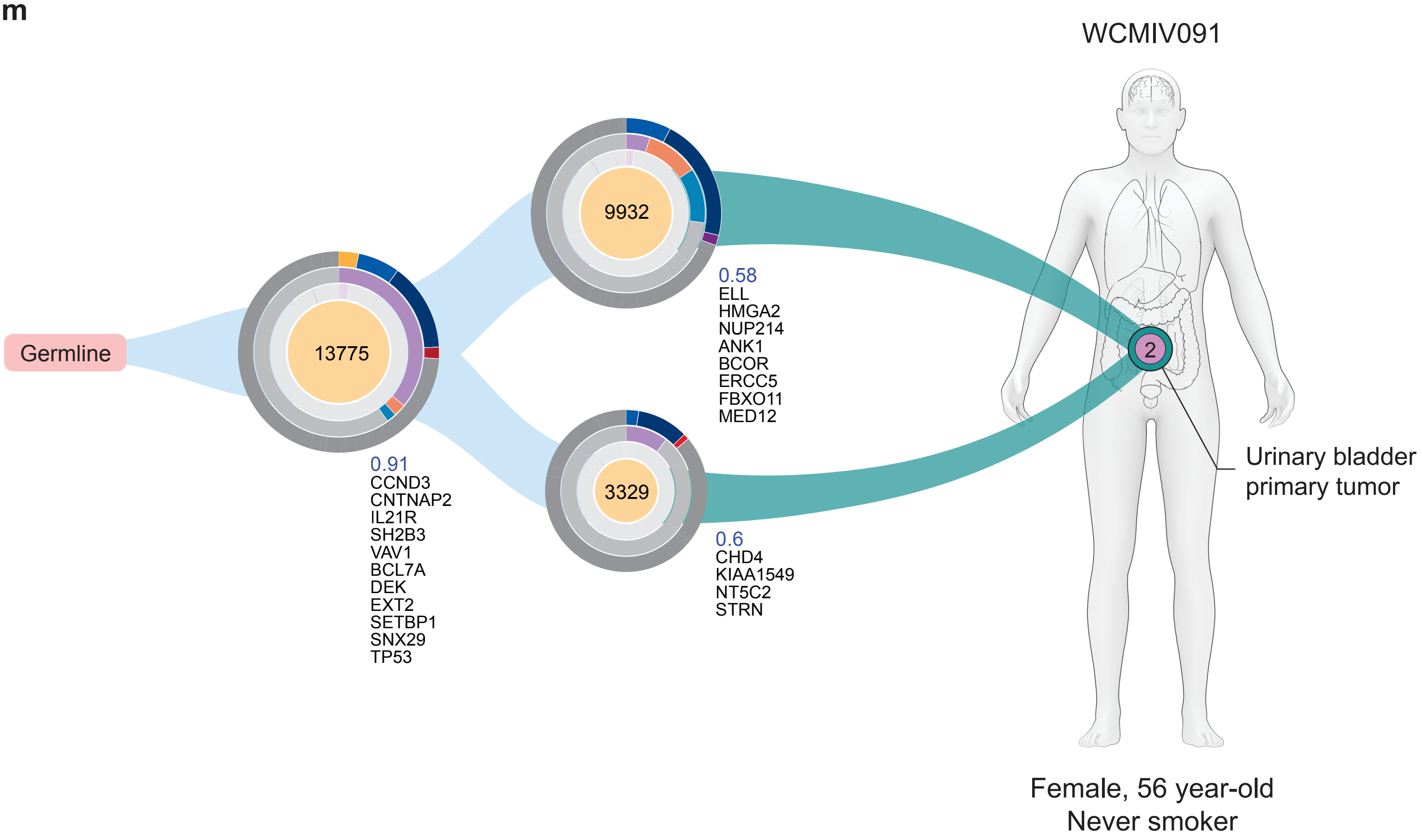

### Supplementary Figure 5

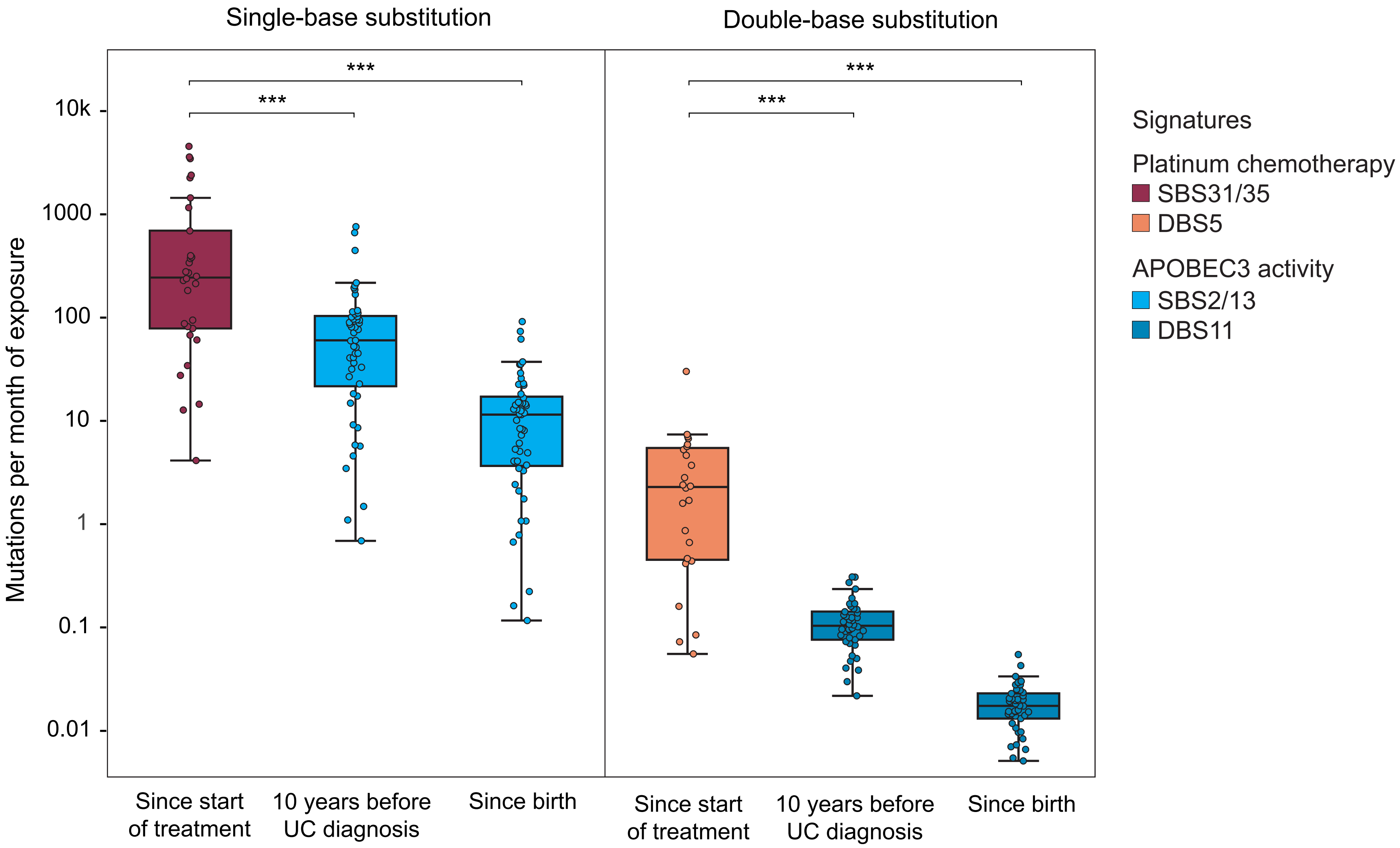


Platinum chemotherapy has higher mutagenic velocity than APOBEC3

### Supplementary Figure 6

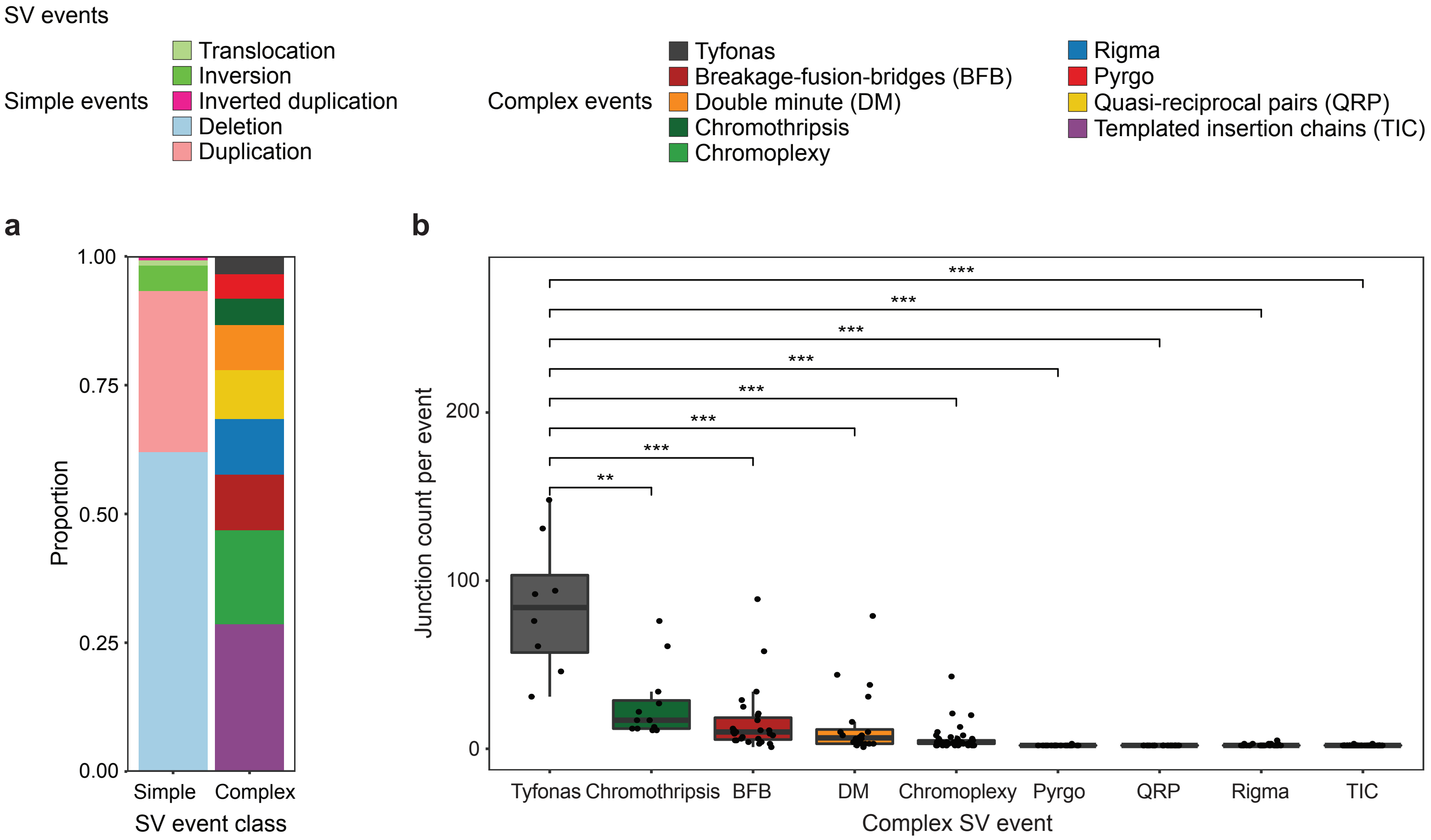

### Supplementary Figure 7

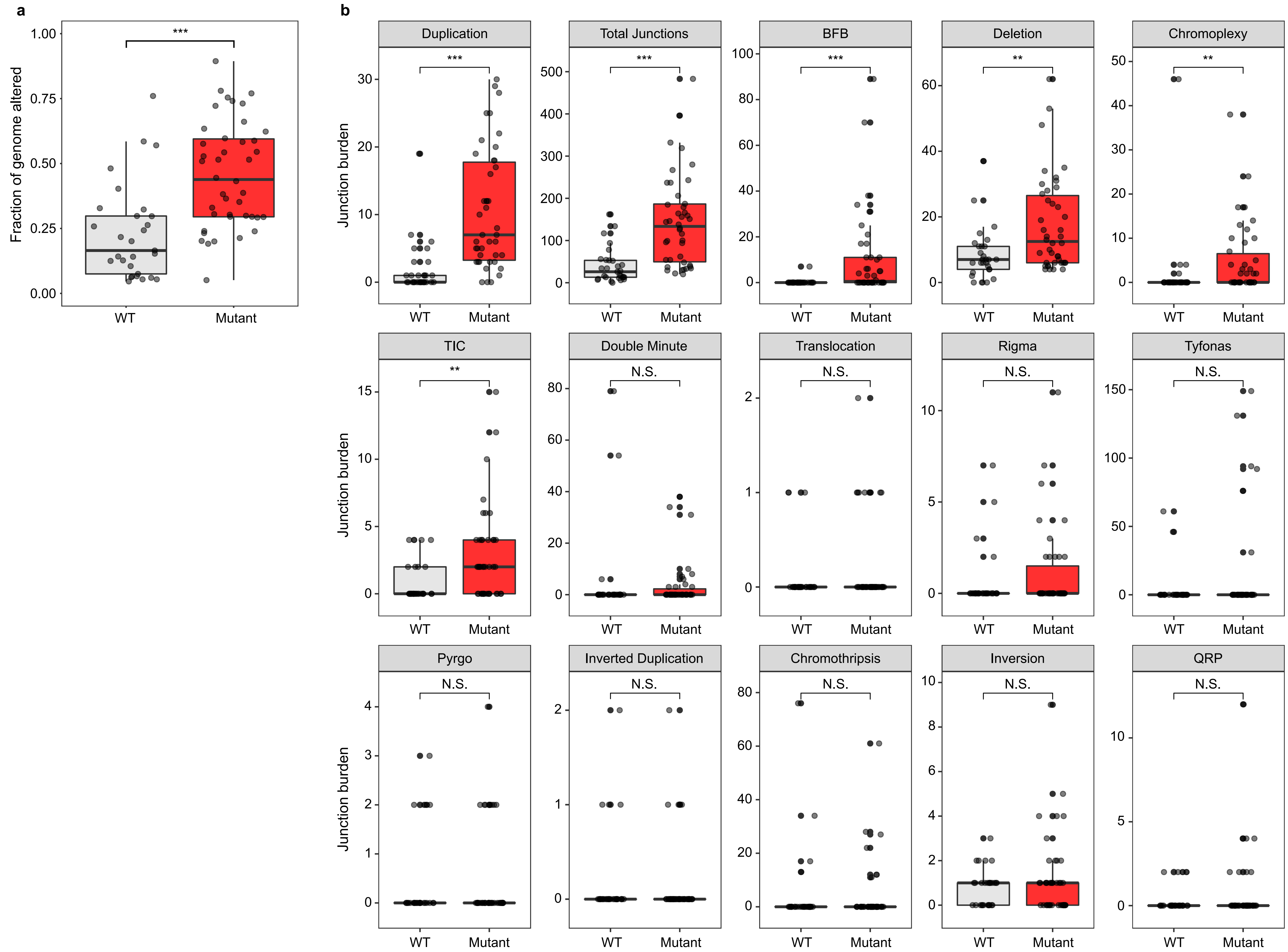

### Supplementary Figure 8

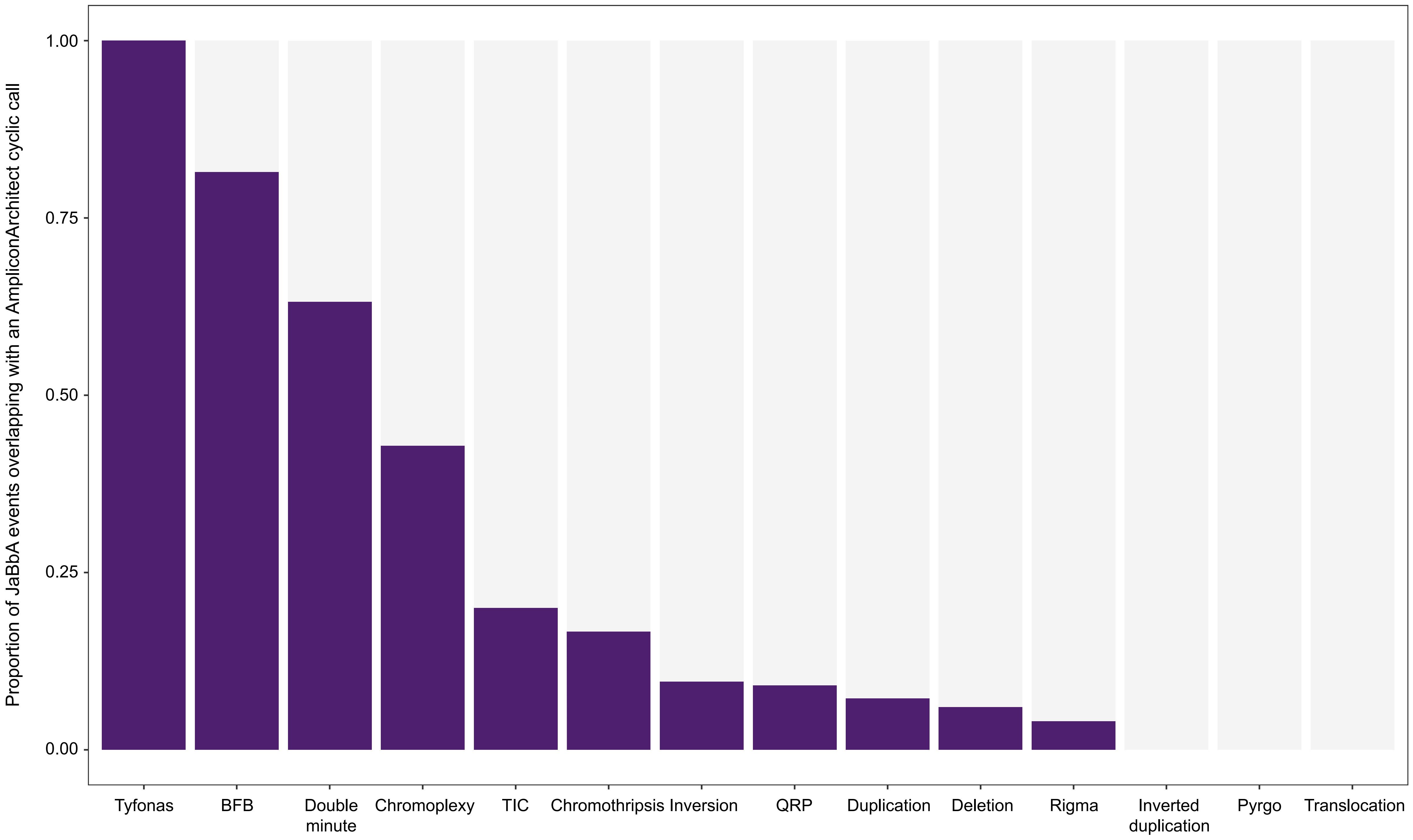

### Supplementary Figure 9a

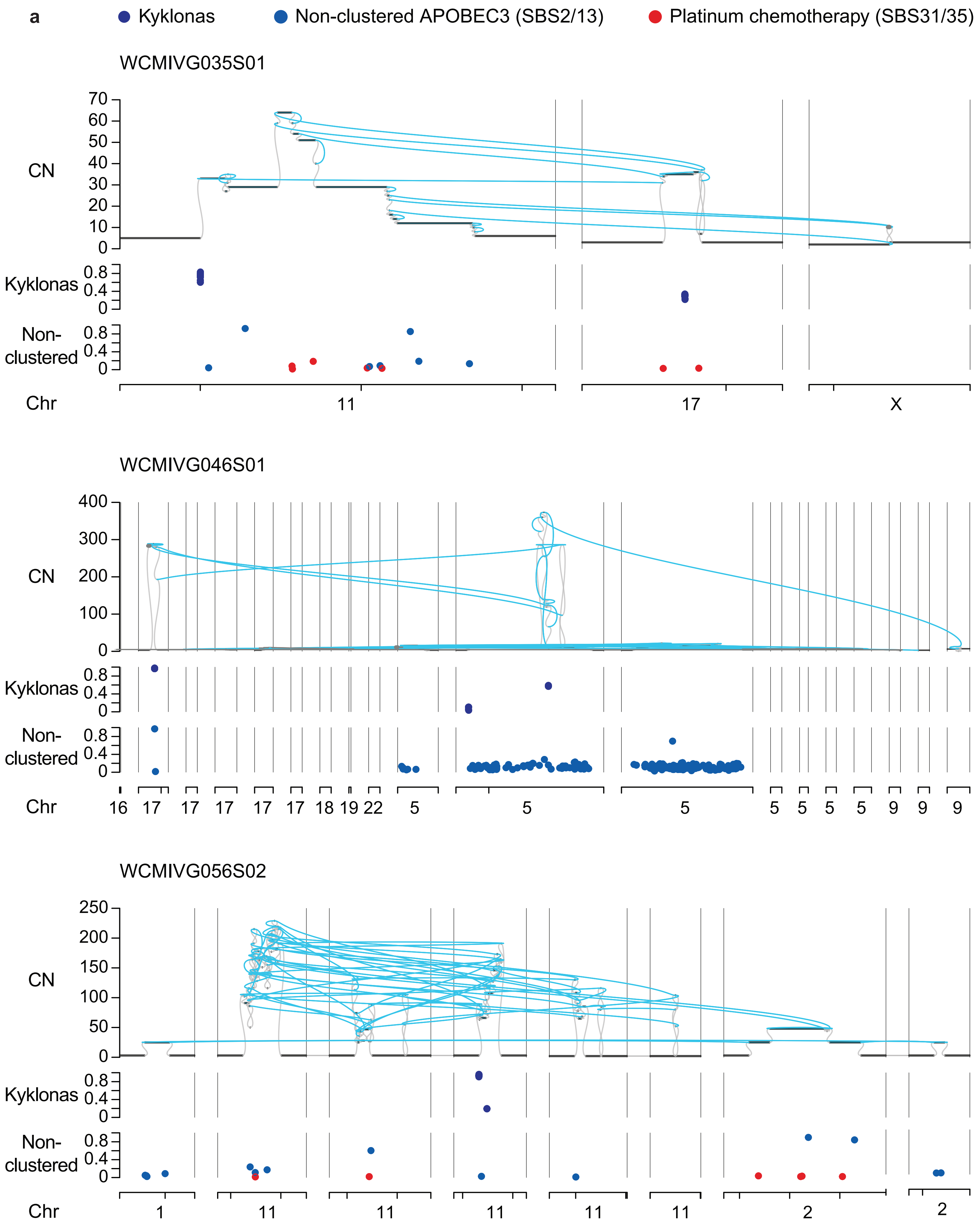

### Supplementary Figure 9b

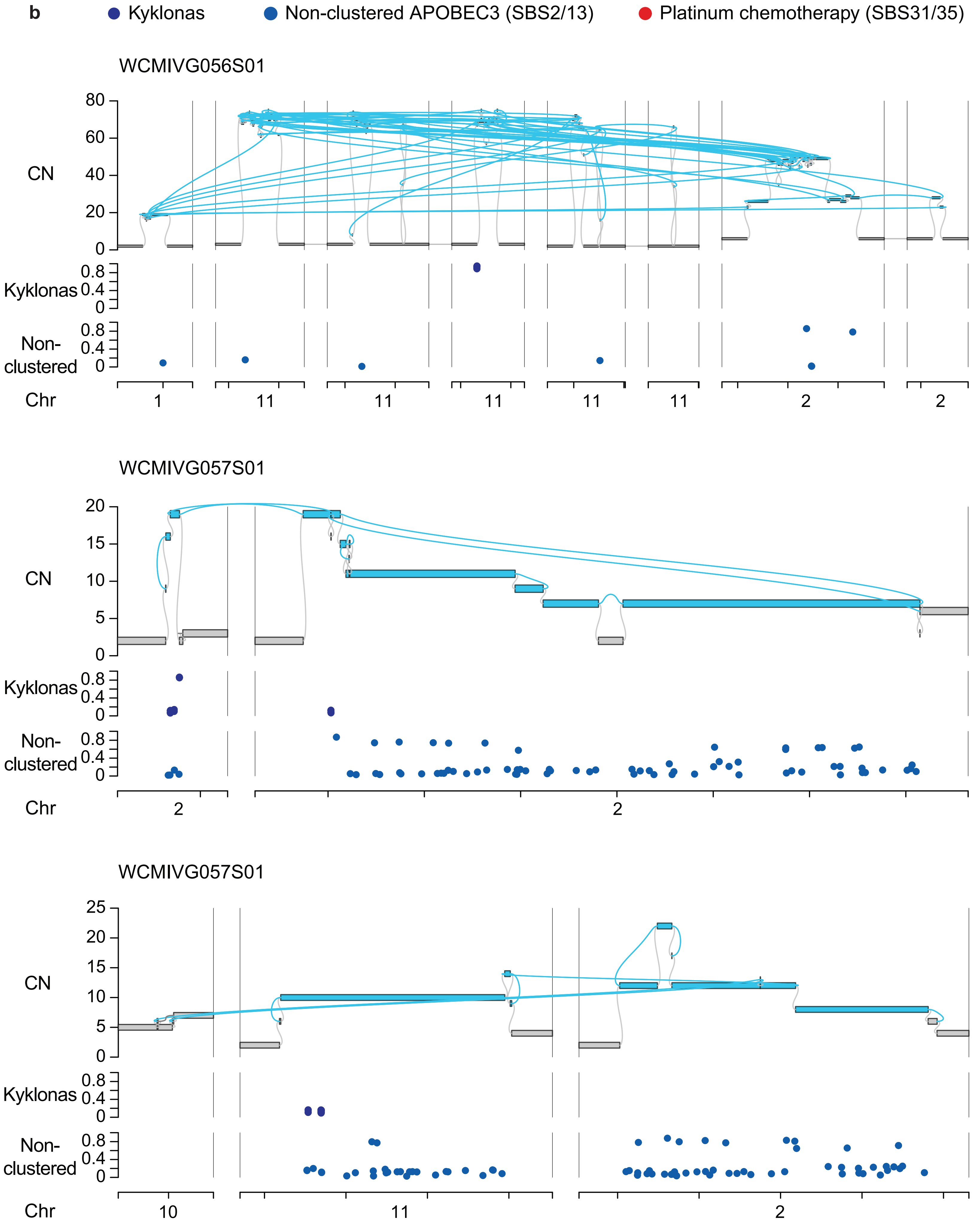

### Supplementary Figure 9c

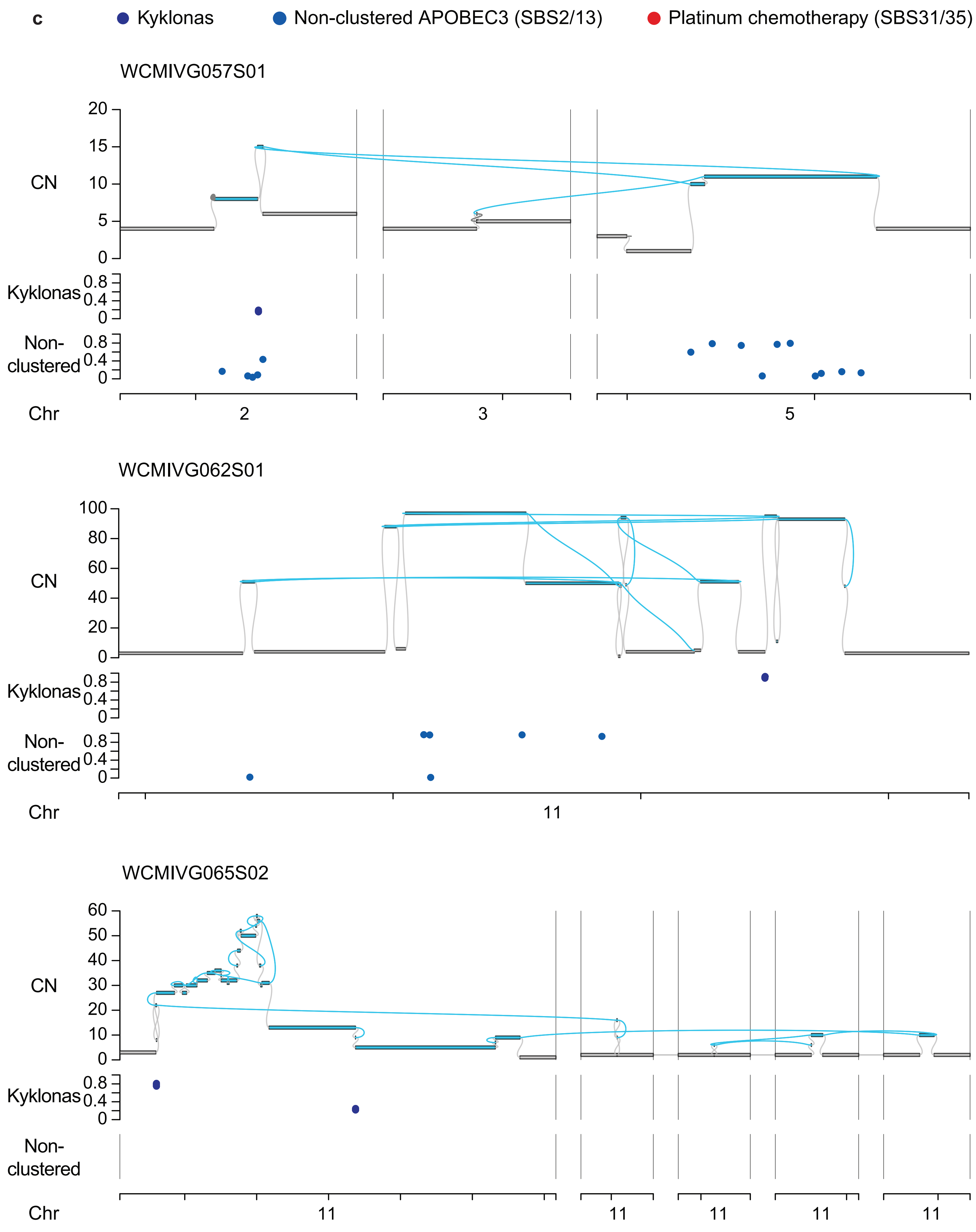

### Supplementary Figure 9d

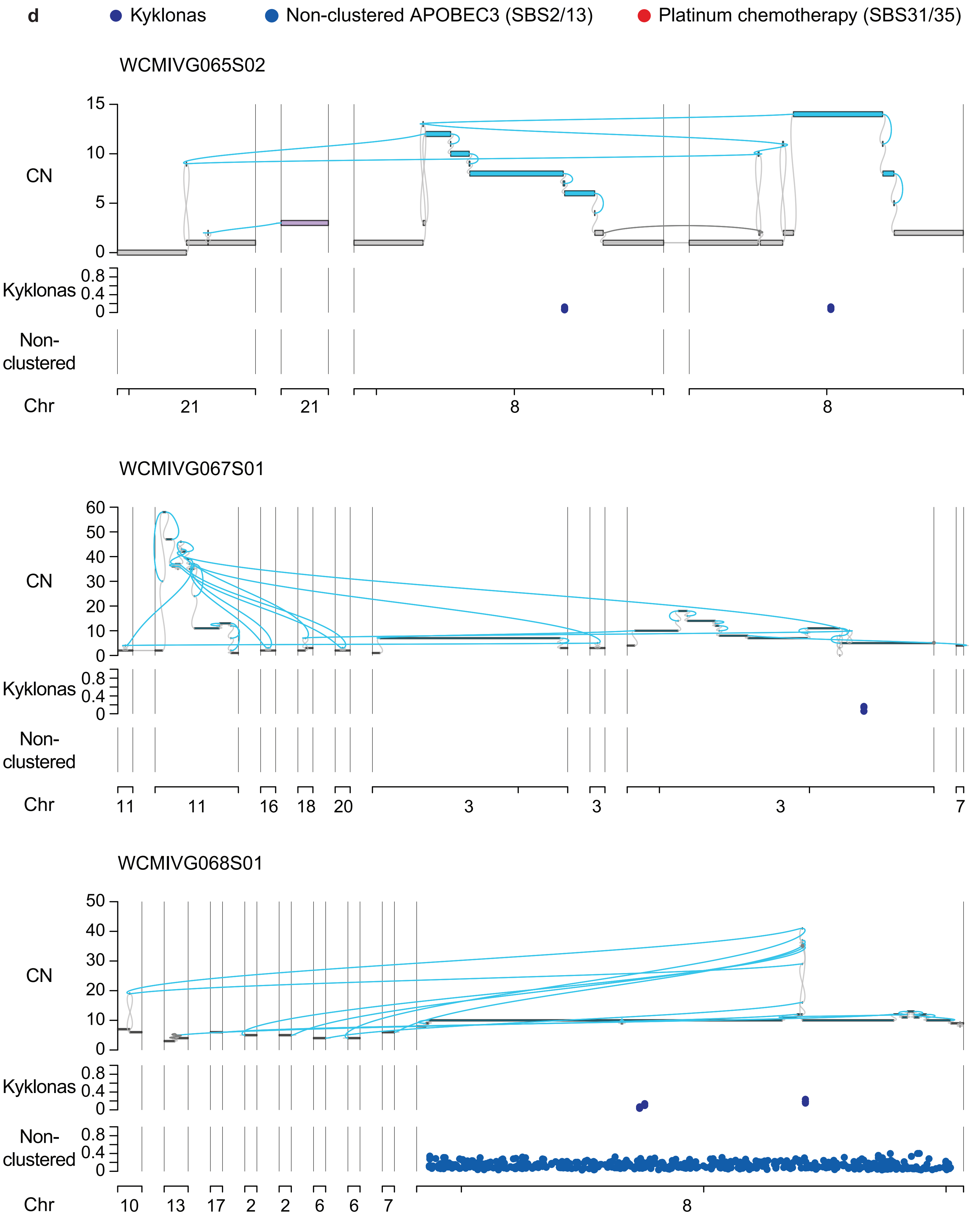

### Supplementary Figure 9e

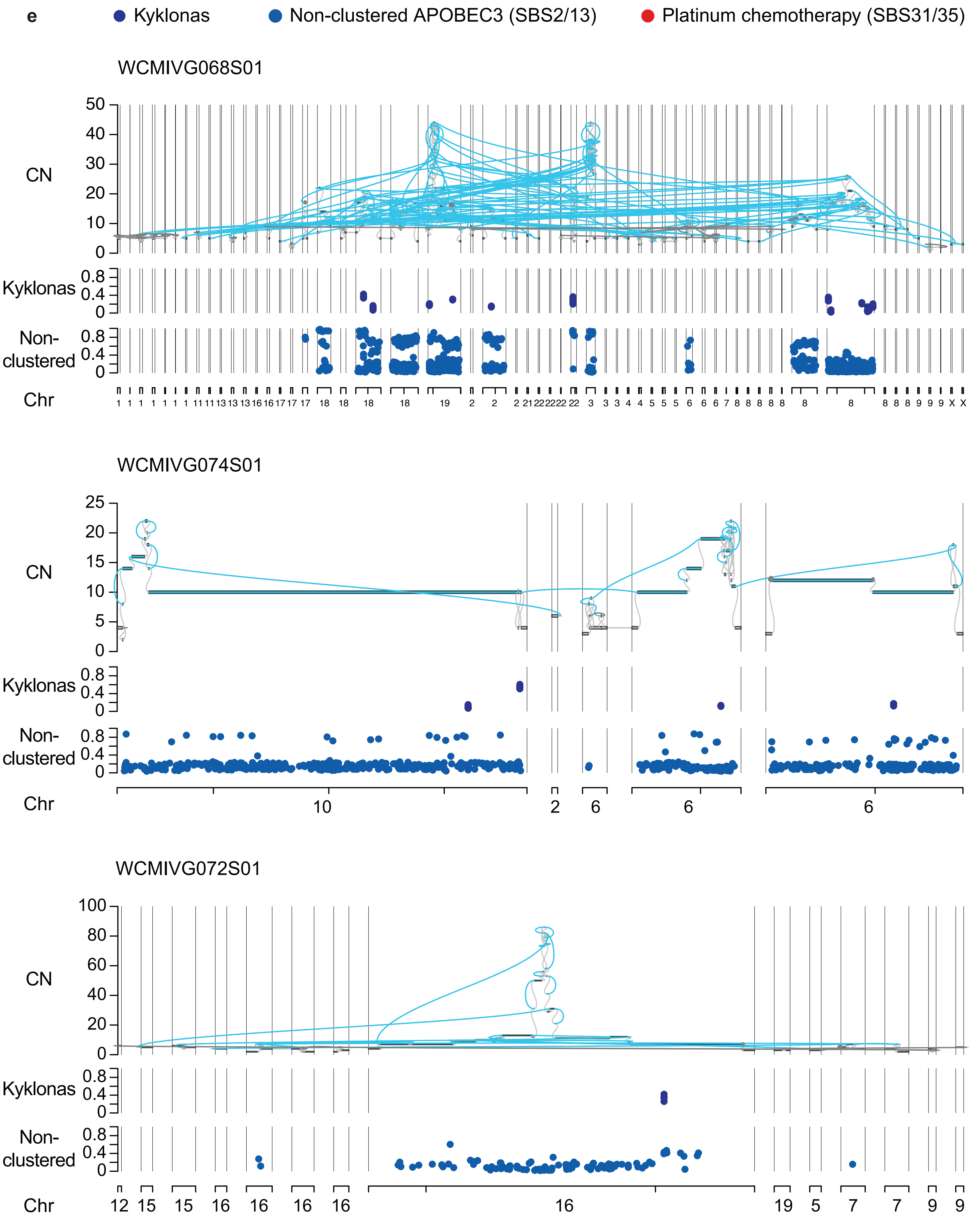

### Supplementary Figure 9f

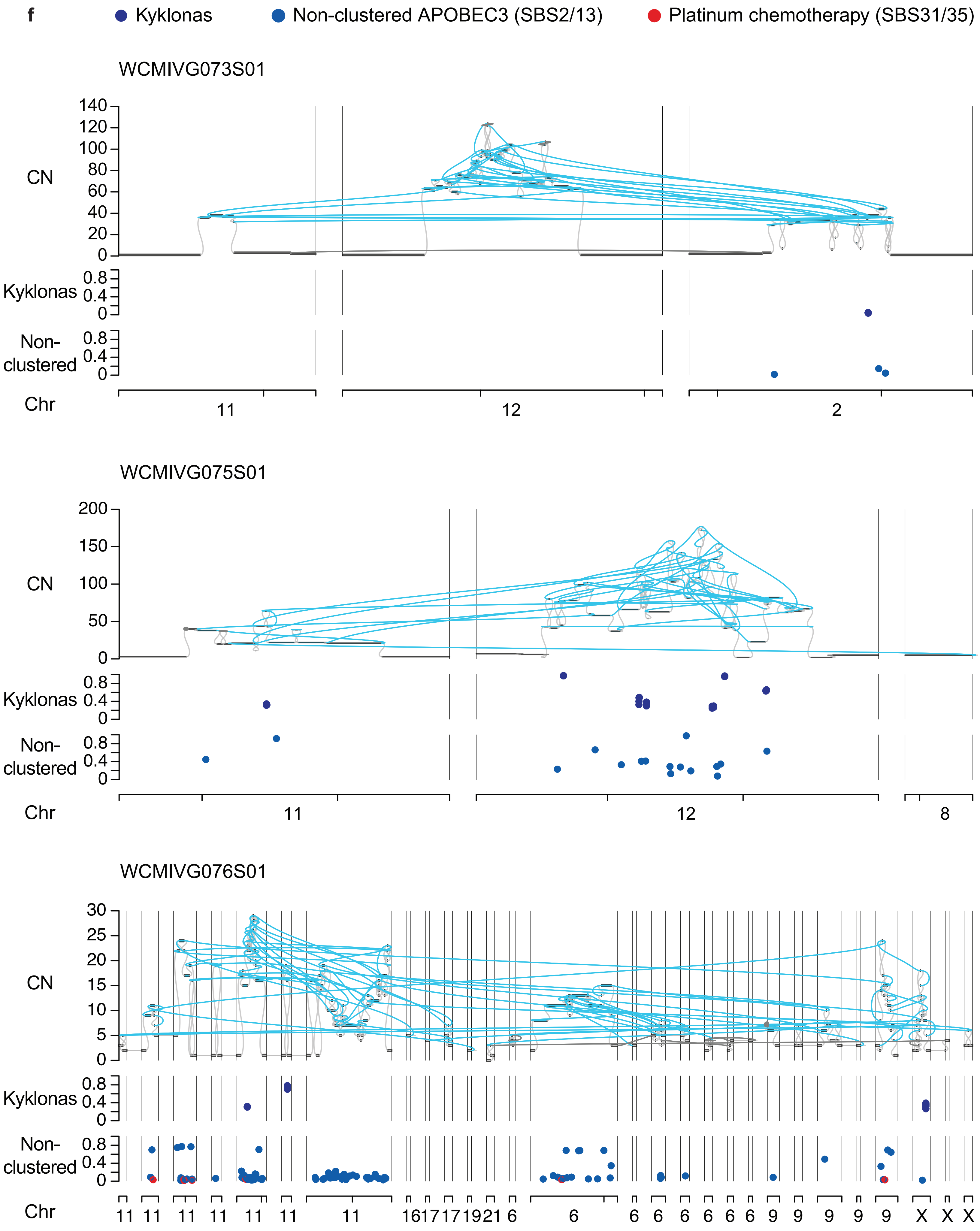

### Supplementary Figure 9g

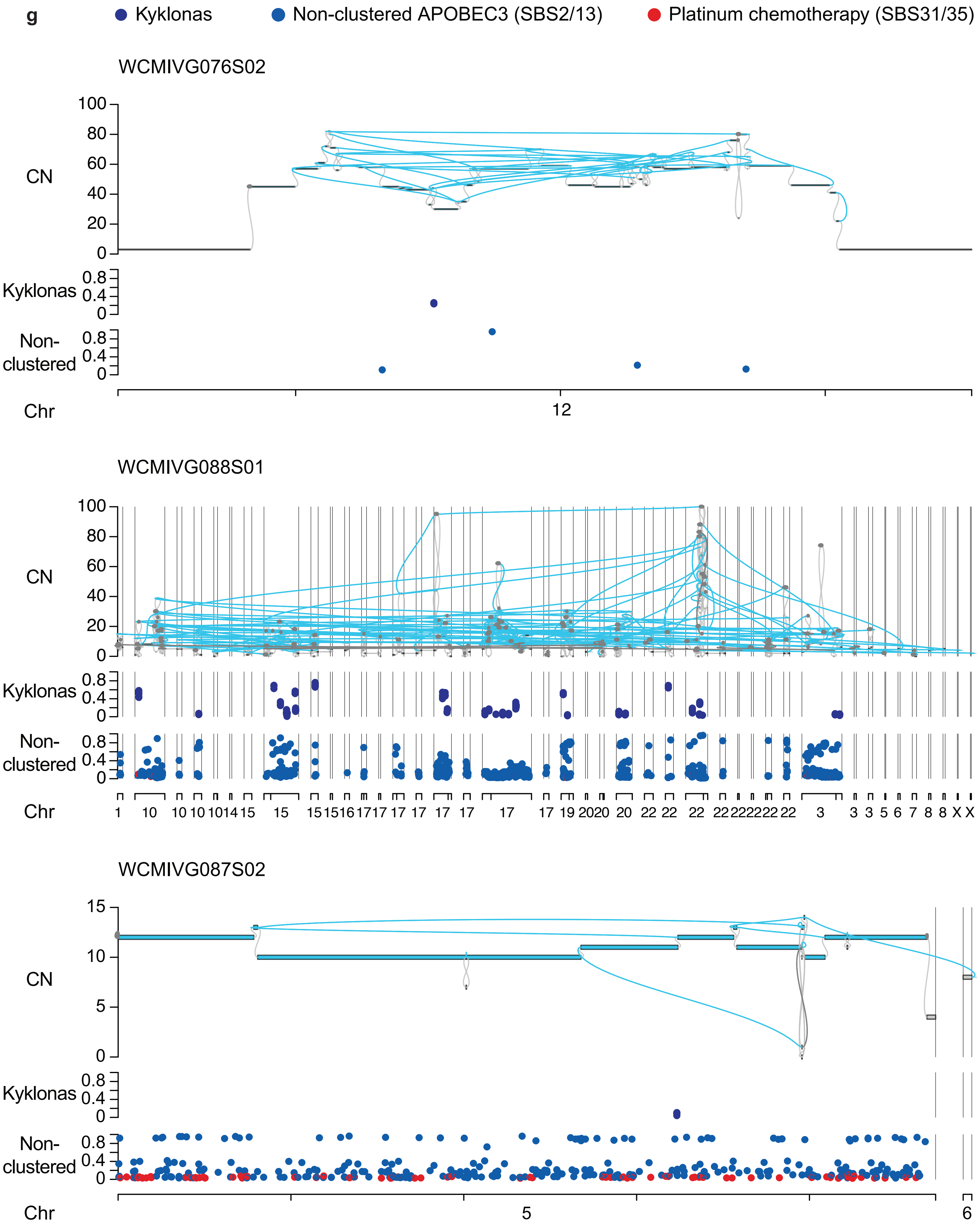

### Supplementary Figure 9h

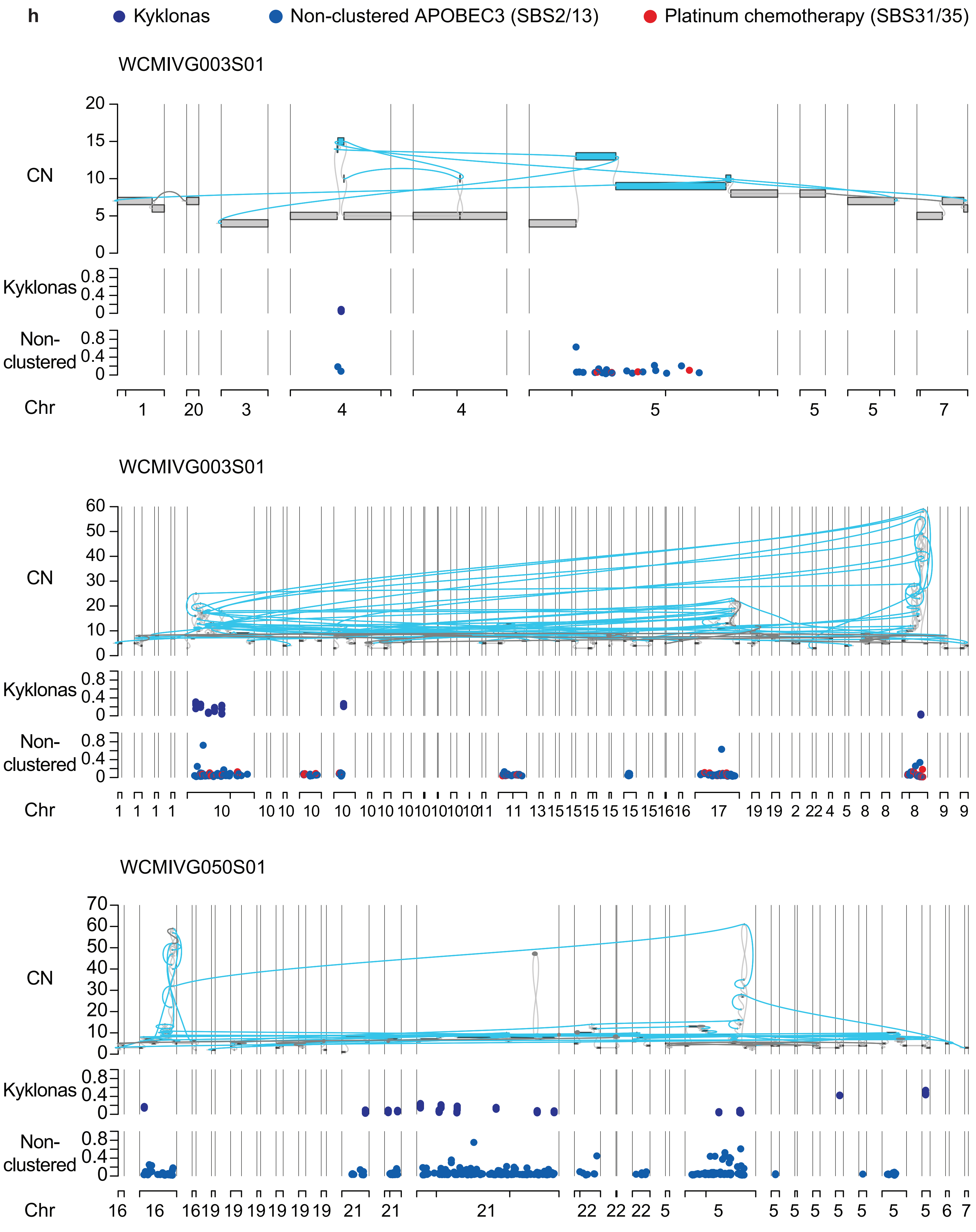

### Supplementary Figure 9i

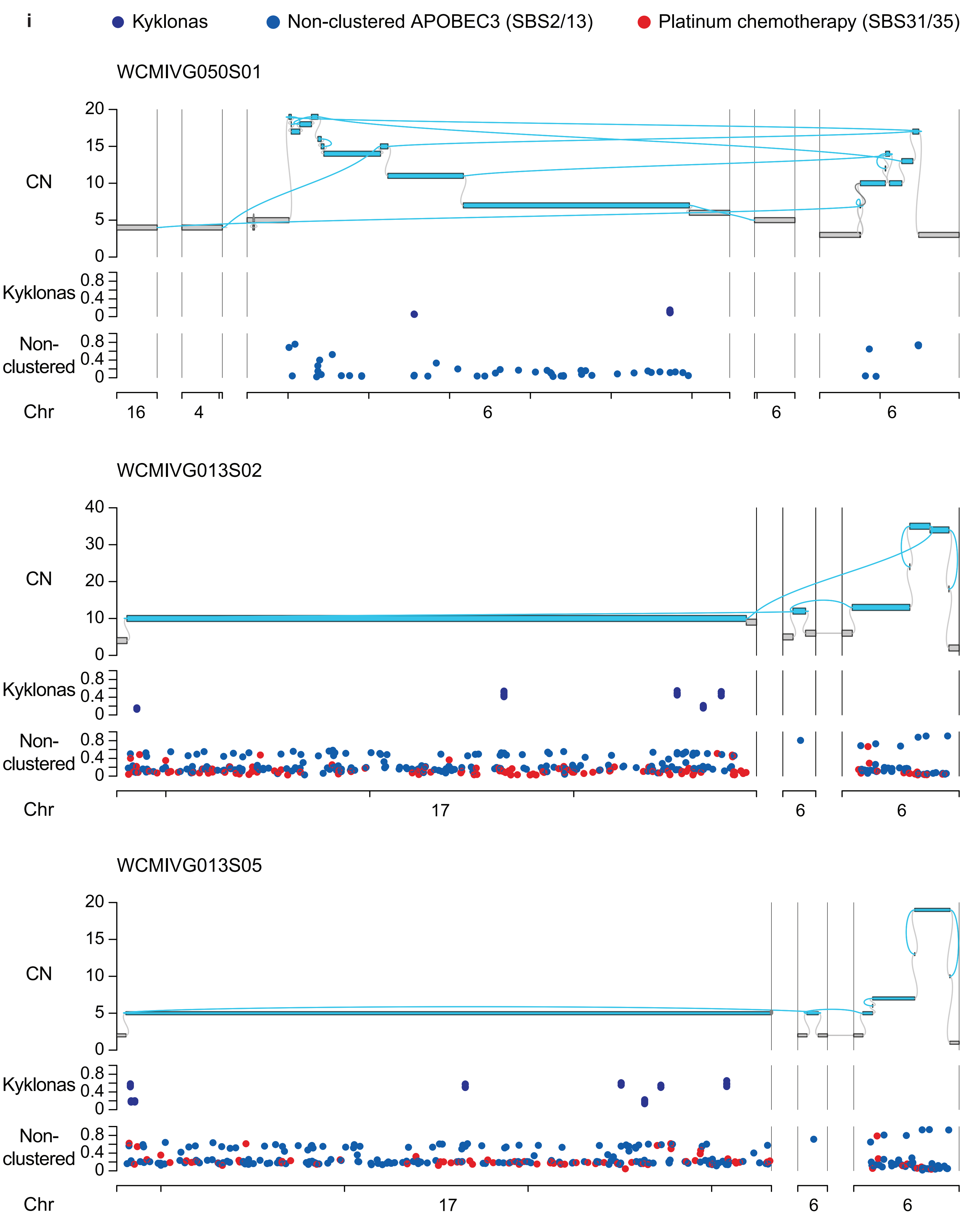

### Supplementary Figure 9j

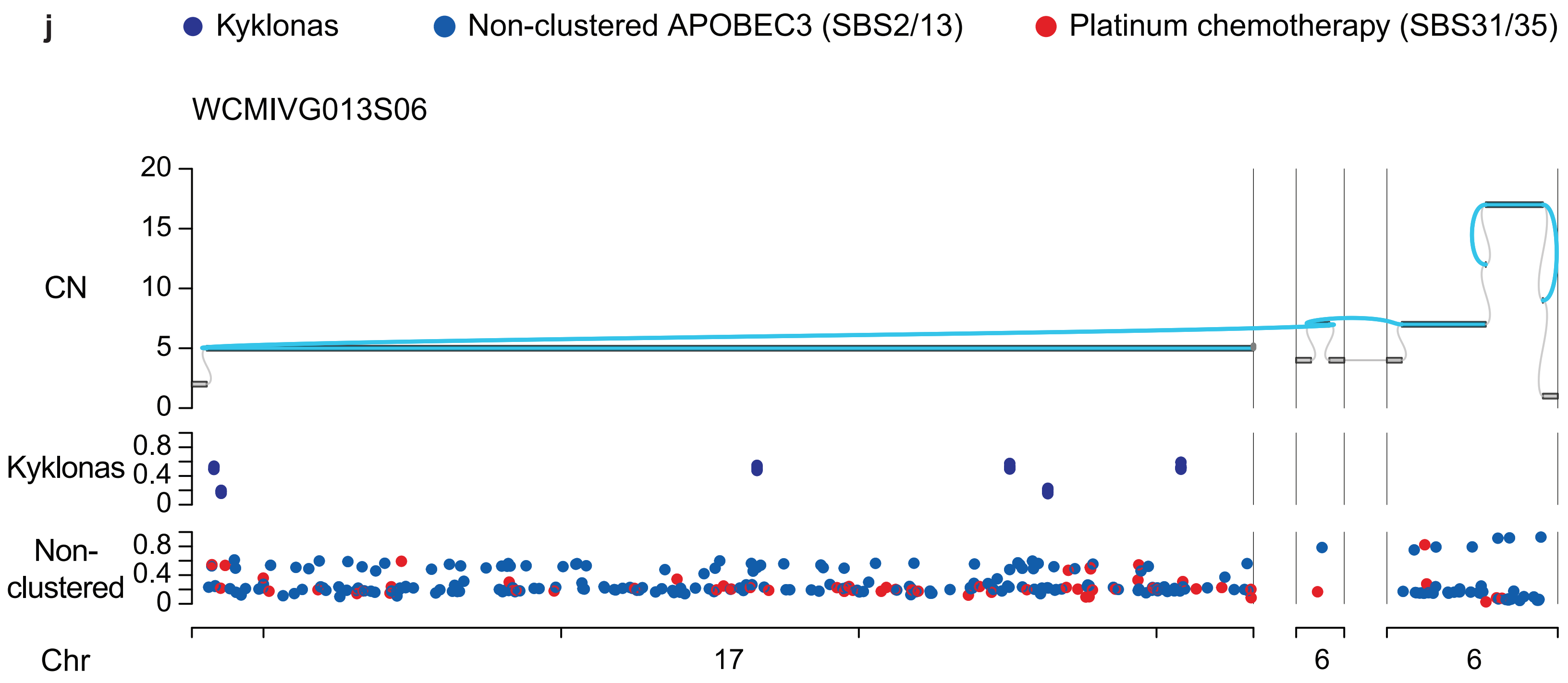
